# Cancer-associated nucleolar morphological changes coincide with altered subcompartment organization and accelerated nascent rRNA trafficking in breast cancer cells

**DOI:** 10.64898/2026.09.22.753621

**Authors:** Kaidi Fan, Wenhao Wang, Gideon Kipkemoi, Yangdulin Li, Yutaka Kikuchi, Haruko Takahashi

**Affiliations:** Graduate School of Integrated Sciences for Life, Hiroshima University, Kagamiyama 1-3-1, Higashi-Hiroshima, Hiroshima, 739-8526 Japan

**Keywords:** Breast cancer, Nucleolus, Ribosome biogenesis, Nucleolar morphology, Nucleolar subcompartments, Nascent rRNA trafficking, Biomolecular condensates

## Abstract

Nucleolar alterations are characteristic features of cancer cells; however, how changes in the overall morphology relate to internal subcompartment organization and ribosomal RNA (rRNA) dynamics remains unclear. In this study, the nucleolar organization of non-transformed MCF-10A mammary epithelial cells and MDA-MB-231 breast cancer cells was quantitatively compared. MDA-MB-231 cells exhibited differences in nucleolar morphology, accompanied by altered spatial organization of the dense fibrillar component and granular component, and accelerated intranucleolar trafficking of nascent rRNA. Pharmacological perturbation with SU056 shifted multiple structural and dynamic parameters toward an MCF-10A-like profile. Conversely, transient physicochemical perturbation with 1,6-hexanediol induced several MDA-MB-231-like nucleolar features in MCF-10A cells, which returned to their untreated profile after washout. These findings support the view that cancer-associated nucleolar alterations are an integrated reorganization across morphological, spatial, and dynamic dimensions. The observed reversibility further indicates that cancer-associated nucleolar organization can be remodeled in response to perturbations. By linking the morphology, subcompartment organization, and nascent rRNA dynamics within a common framework, this study provides a basis for exploring the establishment, maintenance, and remodeling of distinct nucleolar profiles in cancer.

## INTRODUCTION

The sustained growth and proliferation of cancer cells impose high demands on protein synthesis. Accordingly, cancer cells frequently enhance ribosome biogenesis, a nucleolus-centered process that coordinates ribosomal RNA (rRNA) synthesis and processing with the assembly of ribosomal subunits.^1,2,3^ Increased rRNA transcription and dysregulation of multiple components of the ribosome biogenesis machinery have been reported across cancers, including breast cancer.^4,5,6^ Enhanced ribosome biogenesis in cancer is closely associated with changes in nucleolar morphology, particularly nucleolar enlargement, which has been linked to increased rRNA synthesis and proliferative activity.^2,7,8^ Furthermore, cancer cells can exhibit alterations in nucleolar number and shape, and such morphological features have long been used in the pathological assessment of malignancy.^7,9^

While nucleolar size, number, and shape describe its overall morphology, the nucleolus also has a distinct internal architecture that is closely related to its function.^10^ The nucleolus is a nuclear biomolecular condensate organized into three major subcompartments: the fibrillar center (FC), dense fibrillar component (DFC), and granular component (GC).^10,11^ These subcompartments spatially organize successive steps of ribosome biogenesis, with rRNA transcription occurring at the FC–DFC interface, early processing and modification in the DFC, and later processing and ribosomal subunit assembly in the surrounding GC.^10^ Thus, nucleolar organization encompasses the overall morphology and spatial arrangement of functionally distinct internal subcompartments. As a membraneless organelle, the nucleolus maintains distinct internal compartments through molecular interactions rather than physical membrane boundaries.^12^ Liquid–liquid phase separation contributes to this organization through multivalent interactions among proteins and nucleic acids that promote their condensation into phases with distinct compositions and properties.^11,12^ Biophysical studies have shown that the nucleolar subcompartments form coexisting liquid-like phases with distinct molecular compositions and material properties.^10,13^ Differences in intermolecular interactions and surface tensions among these phases contribute to their spatial arrangement, providing a physical basis for the characteristic multiphase organization of the nucleolus.^10,13^ Biomolecular condensates are dynamic assemblies in which molecular components continuously exchange with the surrounding environment, allowing their organization and material properties to respond to changes in molecular interactions and composition.^14^ This dynamic nature suggests that nucleolar organization is capable of remodeling in response to cellular or physicochemical perturbations.^15,16^

Two very recent studies have provided new insights into the dynamic relationship between nucleolar architecture and rRNA processing.^17,18^ Spatial and temporal mapping of pre-rRNA revealed that newly synthesized rRNA undergoes ordered movement from the inner toward the outer regions of the nucleolus as processing proceeds, linking successive stages of rRNA maturation to distinct nucleolar subcompartments.^17,18^ Moreover, perturbation of pre-rRNA processing altered its spatial distribution and nucleolar organization, indicating that rRNA processing and trafficking themselves contribute to the formation and maintenance of nucleolar architecture.^18^ Furthermore, reconstruction of synthetic nucleoli within the nucleus demonstrated that nucleolar architecture depends not only on the organization of molecular components within individual subcompartments but also on pre-rRNA processing and spatial trafficking.^18^ Collectively, these findings highlight a close relationship between nucleolar structure, rRNA processing, and intranucleolar dynamics, providing a more integrated view of nucleolar organization than one based solely on morphology or rRNA transcription.

These recent advances^17,18^ suggest that cancer-associated nucleolar alterations may involve multiple aspects of nucleolar organization. In cancer, alterations have been reported not only in nucleolar morphology and rRNA transcription, but also in multiple molecular pathways involved in rRNA processing, ribosome biogenesis, and the regulation of nucleolar components.^2,3,8^ These observations raise the possibility that the diverse nucleolar alterations described in cancer may occur in a coordinated manner rather than representing entirely independent features associated with malignancy. However, these features have largely been investigated separately, and whether cancer-associated changes in nucleolar morphology are accompanied by alterations in subnucleolar organization and nascent rRNA dynamics remains unclear. Furthermore, whether these different aspects of nucleolar organization can shift together in response to perturbations remains poorly understood. Therefore, integrating these features within the same experimental framework may reveal a broader reorganization of the nucleolus in cancer that cannot be captured by any single morphological or molecular parameter.

In this study, we investigated whether cancer-associated differences in nucleolar morphology are accompanied by changes in the internal subcompartment organization and nascent rRNA dynamics. Non-transformed MCF-10A mammary epithelial cells were compared with MDA-MB-231 triple-negative breast cancer cells, and nucleolar morphology, DFC–GC spatial organization, and intranucleolar trafficking of nascent rRNA were quantitatively analyzed. We further examined how these features respond to pharmacological perturbation in MDA-MB-231 cells and whether cancer-associated nucleolar features could be induced and reversed by transient physicochemical perturbation in MCF-10A cells. These analyses provide an integrated examination of cancer-associated nucleolar organization across the morphological, spatial, and dynamic dimensions.

## RESULTS

### MDA-MB-231 breast cancer cells exhibit distinct changes in nucleolar morphology and internal organization

To characterize cancer-associated changes in nucleolar morphology, non-transformed MCF-10A mammary epithelial cells were compared with MDA-MB-231 cells, an aggressive triple-negative breast cancer cell model. Nucleoli were visualized using immunofluorescence staining of nucleophosmin 1 (NPM1), together with Hoechst 33342 staining of nuclei, and quantitatively analyzed using an automated image-analysis pipeline (see STAR Methods) (Fig. 1A). The MDA-MB-231 cells contained fewer nucleoli per cell than the MCF-10A cells (Fig. 1B). Next, the nucleolar occupancy within the nucleus was quantified by calculating the area of each individual nucleolus relative to the nuclear area, as well as the summed area of all nucleoli within each nucleus relative to the nuclear area. Both measurements were significantly higher in MDA-MB- 231 cells (Fig. 1C and 1D). Consistent with the increased nucleolar area, estimated individual nucleolar volume, calculated from the projected area assuming spherical geometry, was significantly higher in MDA-MB-231 cells (Fig. 1E).

**Figure 1.**
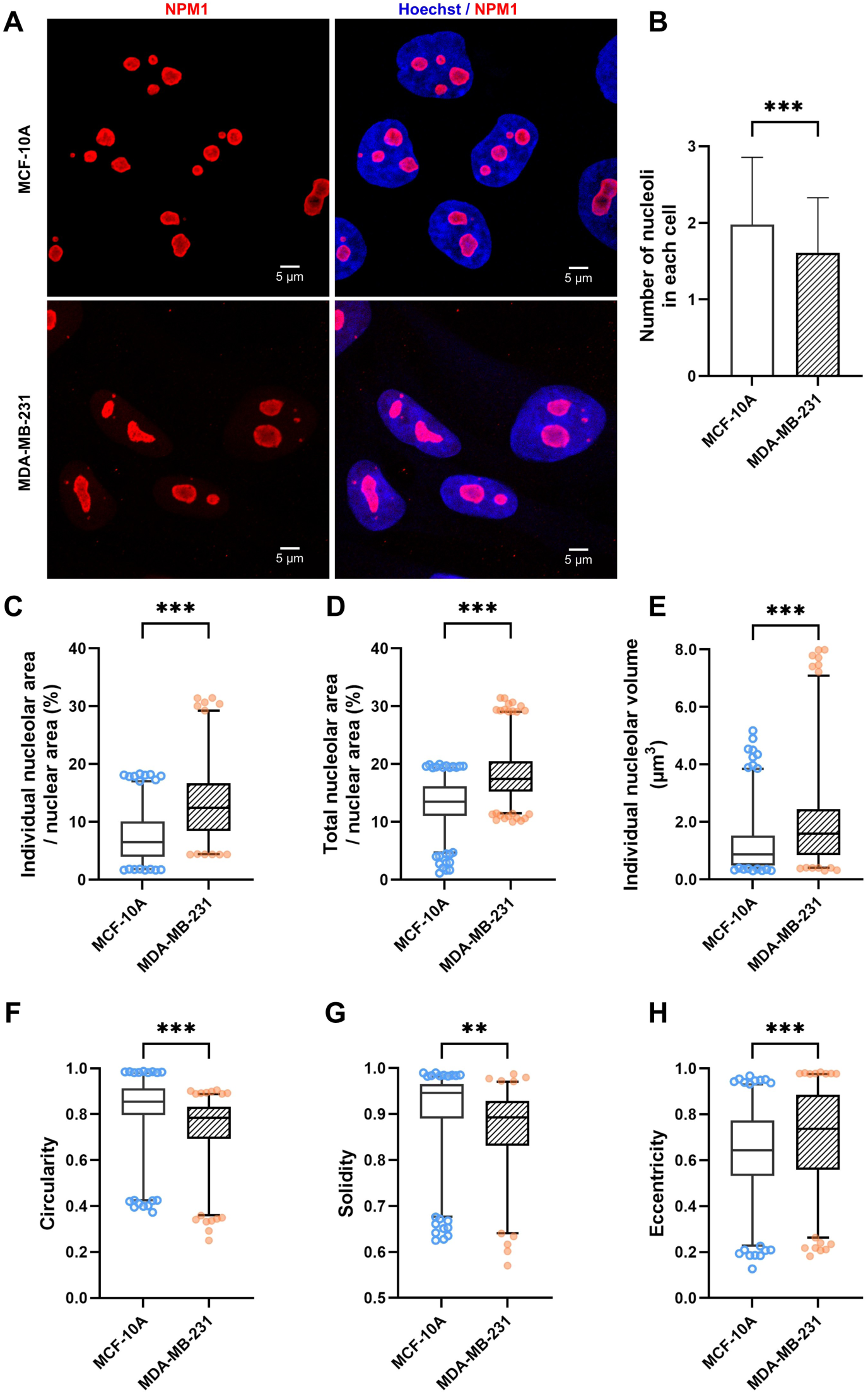
Distinct changes in nucleolar size and shape in MDA-MB-231 breast cancer cells. (A) Representative immunofluorescence images showing nucleolar morphology in normal mammary epithelial cells (MCF-10A), and triple-negative breast cancer cells (MDA-MB-231). Nucleolar marker protein nucleophosmin 1 (NPM1) is shown in red, and nuclei are counterstained with Hoechst 33342 (blue). Scale bar, 5 μm. (B) Distribution of nucleolar number per cell: distribution plots illustrate the number of nucleoli contained within individual cells for each cell line. (C–D) Box-and-whisker plots representing the percentage of (C) single and (D) total nucleolar area per cell. (E–H) Quantitative analysis of nucleolar physical morphology parameters: (E) Estimated volume of individual nucleoli (μm³), calculated from the projected area assuming spherical geometry, (F) Circularity, which indicates the similarity of nucleolar shape to an ideal circle (Data are presented as median with interquartile range), (G) Solidity, which reflects the degree of boundary irregularity, with lower values indicating increased shape concavity, and (H) Eccentricity: values closer to 0 indicate a more spherical shape, whereas values closer to 1 indicate an elongated or elliptical morphology. In total, 950 nucleoli from 492 MCF-10A cells and 821 nucleoli from 500 MDA-MB-231 cells were analyzed over multiple experimental days. The data are presented as box-and-whisker plots with individual data points. Boxes indicate the median and interquartile range, and whiskers indicate the 1st–99th percentiles. Statistical significance was determined using Student’s *t*-test. \*\**p* <0.01, \*\*\**p* <0.001.

Next, nucleolar shape was quantified using circularity, solidity, and eccentricity. Compared with MCF-10A cells, MDA-MB-231 cells showed decreased nucleolar circularity and solidity and increased eccentricity (Fig. 1F–1H). These changes indicate a shift toward less rounded and more angulated nucleoli with increased boundary complexity. Together, these analyses demonstrated distinct differences in nucleolar size and shape between non-transformed MCF-10A and MDA-MB-231 breast cancer cells.

Next, to determine whether the differences in nucleolar size and shape were accompanied by changes in the spatial organization of the DFC and GC, two functionally distinct nucleolar regions involved in successive stages of pre-rRNA maturation were examined. The DFC and GC were visualized using fibrillarin (FBL) and NPM1 as markers, respectively (Fig. 2A). In MCF-10A cells, FBL-positive DFC units were distributed within the surrounding NPM1-positive GC, whereas MDA-MB-231 cells showed a visibly different organization with an increased number of FBL-positive units within the enlarged nucleoli (Fig. 2A). Line profile analysis of FBL and NPM1 fluorescence further illustrated the differences in their spatial distributions between the two cell types (Fig. 2B). Notably, in the MDA-MB-231 nucleoli, NPM1 fluorescence remained high at positions corresponding to the FBL-positive DFC regions, indicating a greater spatial overlap of the two signals.

**Figure 2.**
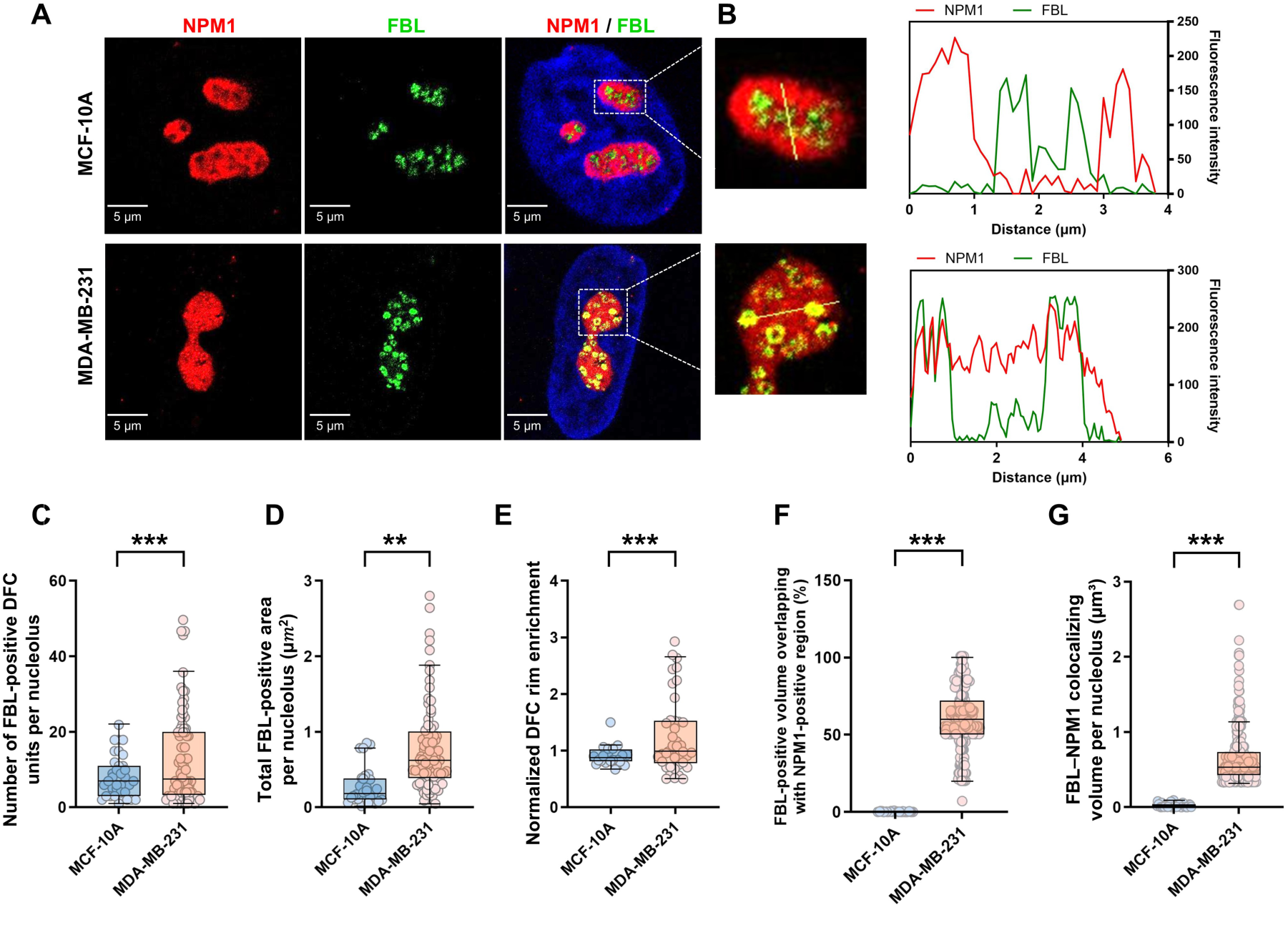
Distinct changes in DFC and GC organization in MDA-MB-231 breast cancer cells. (A) Representative immunofluorescence images showing the spatial distribution of fibrillarin (FBL; green), a marker of the dense fibrillar component (DFC), and nucleophosmin 1 (NPM1)-DsRed (red), a marker of the granular component (GC), in MCF-10A and MDA-MB-231 nucleoli. Scale bar = 5 μm. (B) Representative line profile analysis of FBL (green) and NPM1 (red) fluorescence intensities within individual nucleoli. Yellow lines in the magnified images, corresponding to the dashed boxes in (A), indicate the scan paths used for the intensity profiles shown on the right. (C–D) Quantification of FBL-positive DFC organization, showing (C) the number of FBL-positive DFC units per nucleolus and (D) the total FBL-positive area per nucleolus. (E) Quantification of the radial distribution of FBL-positive DFC units using normalized DFC rim enrichment. A value of 1 represents a uniform radial distribution, whereas values above or below 1 indicate relative enrichment toward the nucleolar periphery or center, respectively. (F–G) Three-dimensional quantification of FBL–NPM1 spatial overlap: (F) percentage of FBL-positive volume overlapping with the NPM1-positive region per nucleolus and (G) total FBL–NPM1 colocalizing volume per nucleolus. Representative three-dimensional reconstructions are shown in Videos S1 and S2. A total of 770 FBL-positive DFC units from 100 nucleoli in 52 MCF-10A cells and 1,641 FBL-positive DFC units from 89 nucleoli in 54 MDA-MB-231 cells were analyzed over multiple experimental days. The data are presented as box-and-whisker plots with individual data points. Boxes indicate the median and interquartile range, and whiskers indicate the 1st–99th percentiles. Statistical significance was determined using Student’s *t*-test. \*\**p* < 0.01, \*\*\**p* < 0.001.

Quantitative analysis showed that MDA-MB-231 nucleoli contained more FBL-positive DFC units and a larger total FBL-positive area per nucleolus than MCF-10A nucleoli (Fig. 2C and 2D). In both cell types, the number of FBL-positive DFC units was positively associated with nucleolar area (Fig. S1 in Supplementary Information (SI)). Individual DFC units also differed in estimated volume and shape between MCF-10A and MDA-MB-231 cells (Fig. S2). Next, the radial distribution of DFC units was assessed using normalized DFC rim enrichment, in which a value of 1 represents a uniform distribution, and higher values indicate greater enrichment toward the nucleolar periphery. DFC rim enrichment significantly increased in MDA-MB-231 cells (Fig. 2E), indicating that DFC units were preferentially distributed closer to the nucleolar periphery toward the outer region of the surrounding GC. Consistent with the line profile analysis (Fig. 2B), three-dimensional colocalization analysis showed an increased percentage of FBL-positive volume overlapping with the NPM1-positive region (Fig. 2F), together with an increase in the total FBL–NPM1 colocalizing volume (Fig. 2G; Videos S1 and S2). Together, these results demonstrate that differences in nucleolar size and shape in MDA-MB-231 cells are accompanied by changes in the spatial organization of DFC and GC.

### Accelerated intranucleolar trafficking of nascent rRNA in breast cancer cells

Next, whether the differences in DFC–GC organization were accompanied by changes in the intranucleolar dynamics of nascent rRNA was examined. Nascent rRNA trafficking was quantified using a fluorescence pulse–chase labeling approach (Fig. 3A and 3B). Nascent rRNA was labeled with 5-ethynyl uridine (5-EU) and tracked over time relative to FBL-positive DFC and NPM1-positive GC regions. In MCF-10A cells, the 5-EU signal gradually redistributed from the DFC-associated regions toward the surrounding GC during the chase period. In contrast, MDA-MB-231 cells showed more rapid redistribution of the 5-EU signal toward the GC (Fig. 3A and 3B).

**Figure 3.**
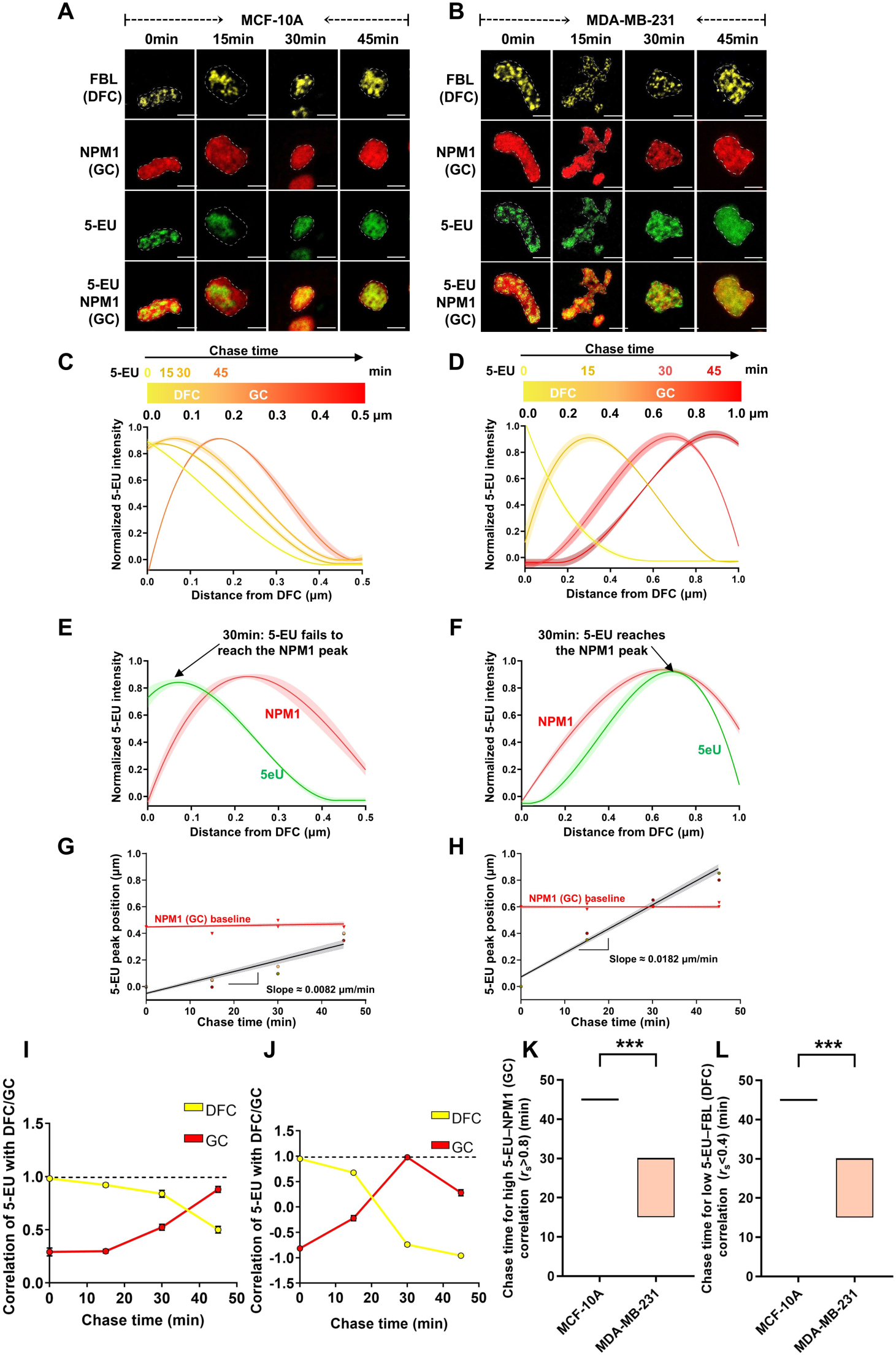
Accelerated intranucleolar trafficking of nascent rRNA in MDA-MB-231 cells. (A–B) Representative fluorescence pulse–chase images showing the intranucleolar spatiotemporal distribution of nascent rRNA in (A) MCF-10A and (B) MDA-MB-231 cells at the indicated chase time points (0, 15, 30, and 45 min). Nascent RNA was pulse-labeled with 5-ethynyl uridine (5-EU; green). Fibrillarin (FBL; yellow) and nucleophosmin 1 (NPM1; red) indicate the dense fibrillar component (DFC) and granular component (GC), respectively. Dashed lines indicate the nucleolar boundaries. Scale bars: 5 μm. (C–D) Spatial profiles of 5-EU fluorescence intensity in (C) MCF-10A and (D) MDA-MB-231 cells during the chase period. The y-axis shows min–max normalized fluorescence intensity, and the x-axis shows the distance from the DFC center (μm). (E–F) Spatial profiles of 5-EU (green) and NPM1 (red) fluorescence intensities after 30 min of chase in (E) MCF-10A and (F) MDA-MB-231 cells. The intensities were min–max normalized and plotted as a function of the distance from the DFC center. (G–H) Estimation of apparent intranucleolar translocation velocity in (G) MCF-10A and (H) MDA-MB-231 cells. The scatter plots show the displacement of the 5-EU signal peak over time, and the linear regression fits are shown in black. The slope of the regression line was used to estimate the apparent translocation velocity. NPM1 peak positions are shown as a reference (red). (I–J) Correlation analysis (Spearman’s correlation coefficient) between the 5-EU signal and FBL or NPM1 over the chase period in (I) MCF-10A and (J) MDA-MB-231 cells. (K) Chase time required to reach a Spearman correlation coefficient *r*ₛ > 0.8 between 5-EU (nascent rRNA) and NPM1 (GC). (L) Chase time required to reach a Spearman correlation coefficient *r*ₛ < 0.4 between 5-EU (nascent rRNA) and FBL (DFC). For (C–F), solid lines and shaded areas represent mean ± standard error of the mean. For (K–L), data are presented as mean ± standard deviation. The numbers of nucleoli analyzed at 0, 15, 30, and 45 min were 60, 59, 67, and 60 for MCF-10A cells (31, 31, 35, and 31 cells, respectively) and 79, 85, 87, and 72 for MDA-MB-231 cells (48, 52, 53, and 44 cells, respectively). Statistical significance between two groups was determined using Student’s *t*-test. \*\*\**p* < 0.001.

Line profile analysis showed that the spatial peak of the 5-EU signal shifted outward more rapidly in MDA-MB-231 cells than in MCF-10A cells (Fig. 3C and 3D). After 30 min of the chase, the 5-EU signal in MDA-MB-231 cells showed greater spatial overlap with the NPM1-positive GC regions, whereas in MCF-10A cells, the signal remained more centrally distributed (Fig. 3E and 3F). Regression analysis of the displacement of the 5-EU signal peak yielded a higher apparent intranucleolar translocation velocity in MDA-MB-231 cells (0.0182 μm/min) than in MCF-10A cells (0.0082 μm/min) (Fig. 3G and 3H).

Further, the temporal relationship between nascent rRNA and the DFC and GC was examined using correlation analysis. The association of the 5-EU signal with FBL decreased over the chase period, whereas its association with NPM1 increased, consistent with the redistribution of nascent rRNA from DFC-associated to GC-associated regions (Fig. 3I and 3J). In MDA-MB-231 cells, the 5-EU–NPM1 correlation reached its maximum at approximately 30 min. Consistent with this earlier redistribution, MDA-MB-231 cells required a shorter chase time to achieve a high correlation between 5-EU and NPM1 and a low correlation between 5-EU and FBL than MCF-10A cells (Fig. 3K and 3L). Together, these results demonstrate that MDA-MB-231 cells exhibit accelerated intranucleolar trafficking of nascent rRNA compared with MCF-10A cells, indicating that the differences in nucleolar morphology and internal organization are accompanied by altered rRNA dynamics.

### SU056 treatment shifts MDA-MB-231 nucleolar features toward an MCF-10A-like profile

Having identified the differences in nucleolar morphology, DFC–GC organization, and nascent rRNA dynamics between MCF-10A and MDA-MB-231 cells, we next examined whether these cancer-associated nucleolar features could be altered by pharmacological perturbation. YB-1 has been implicated in aggressive breast cancer phenotypes, including those of MDA-MB-231 cells.^19^ SU056 has subsequently been reported to target YB-1 and suppress the growth of triple-negative breast cancer cells, including MDA-MB-231 cells, with proteomic analysis revealing broad changes in ribosome- and translation-associated proteins following SU056 treatment.^24^ Therefore, whether the molecular changes associated with SU056 treatment extended to factors involved in nucleolar function and rRNA processing was investigated.

RNA-seq data^20,21,22,23^ comparing MCF-10A and MDA-MB-231 cells were integrated with the published proteomic dataset from SU056-treated MDA-MB-231 cells^24^, focusing on the 331 ribosome biogenesis–related genes defined by Zang *et al.*^25^ (Fig. S3). This analysis identified 19 candidates that were upregulated in MDA-MB-231 cells and decreased following SU056 treatment, as well as 8 candidates that were downregulated in MDA-MB-231 cells and increased following SU056 treatment (Figs. S3A–S3C). These candidates included multiple factors associated with nucleolar function and rRNA processing. Gene Ontology analysis further revealed the enrichment of processes related to ribosome assembly, rRNA modification, and ribosome biogenesis (Fig. S3D).

These *in silico* findings prompted us to examine whether SU056 treatment alters the nucleolar profile of MDA-MB-231 cells. Cell viability and nucleolar morphology were first evaluated across a range of SU056 concentrations (0.1–2.0 μM). Cell viability decreased with increasing SU056 concentration (Fig. S4A), whereas quantitative morphological analysis revealed changes in multiple nucleolar parameters across the concentration range (Figs. S4B–S4H). Based on these analyses, SU056 concentrations of 0.25 and 2.0 μM were selected for subsequent quantitative imaging to represent mild and strong perturbation conditions, respectively.

The nucleolar morphology and internal DFC–GC organization were then examined using the same quantitative parameters used to characterize MCF-10A and DMSO-treated MDA-MB-231 cells (Fig. 4A and 4B). Line profile analysis showed that SU056 treatment increased the spatial separation between the FBL and NPM1 signals, with reduced NPM1 fluorescence at positions corresponding to FBL-positive DFC regions (Fig. 4B). SU056 treatment also reduced individual nucleolar area and increased circularity, shifting both parameters toward the values observed in MCF-10A cells (Fig. 4C and 4D). Additional morphological parameters, including the nucleolar number, total nucleolar area, solidity, and eccentricity, showed corresponding changes toward an MCF-10A-like profile across the SU056 concentration range (Figs. S4C–S4H).

**Figure 4.**
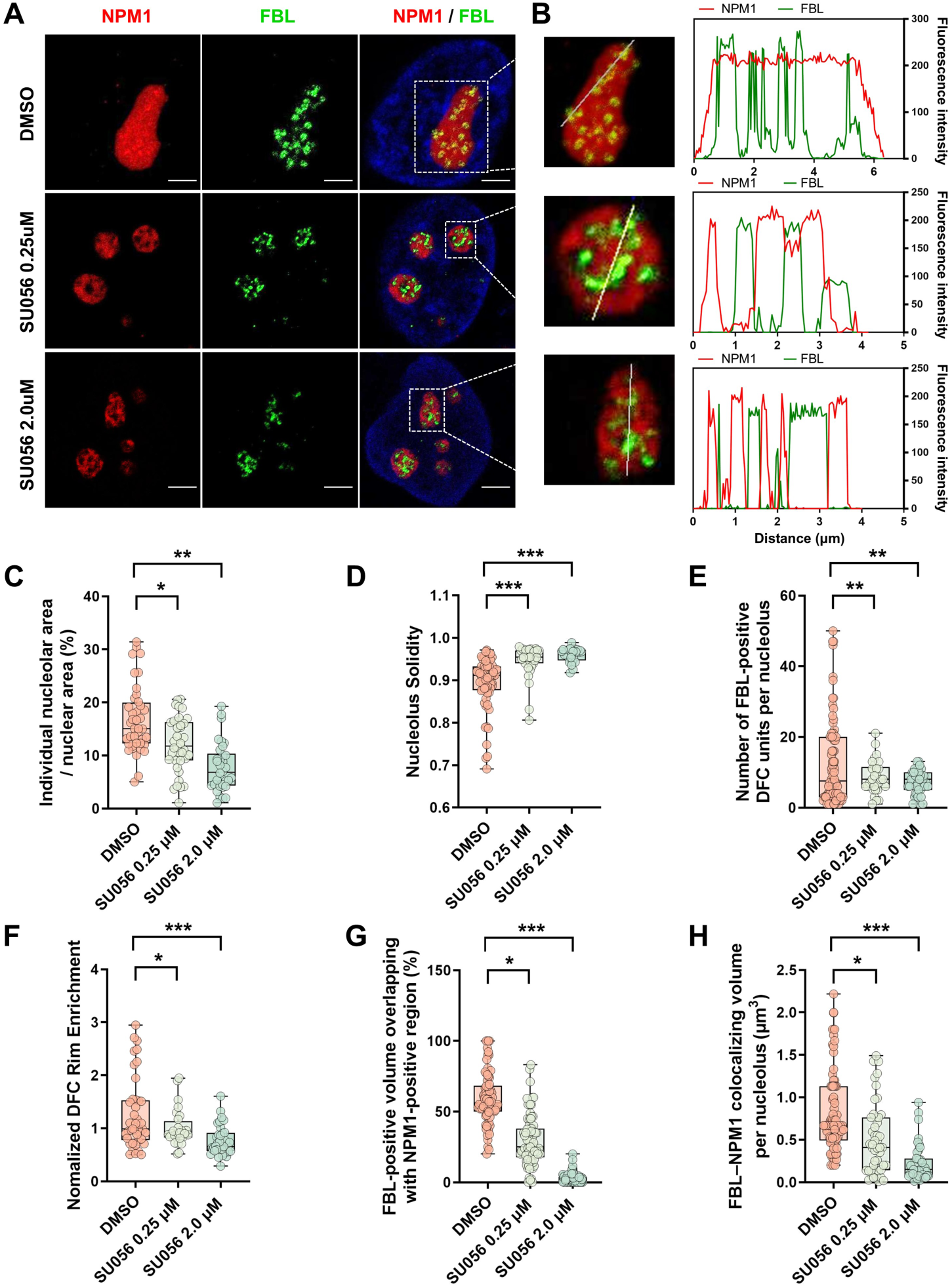
SU056 treatment shifts nucleolar morphology and DFC–GC organization toward an MCF-10A-like profile in MDA-MB-231 cells. (A) Representative immunofluorescence images of nucleophosmin 1 (NPM1; red, granular component [GC]) and fibrillarin (FBL; green, dense fibrillar component [DFC]) in MDA-MB-231 cells treated with dimethyl sulfoxide (DMSO) or SU056 at concentrations of 0.25 and 2.0 μM. Scale bar = 5 μm. (B) Representative line profile analysis of NPM1 (red) and FBL (green) fluorescence intensities. Yellow lines in the magnified insets corresponding to the dashed boxes in (A) indicate the scan paths for the intensity profiles shown on the right. The x-axis represents the distance along the scan line (μm), and the y-axis represents fluorescence intensity. (C– D) Quantification of nucleolar morphology: (C) individual nucleolar area relative to nuclear area and (D) circularity. (E–F) Quantification of DFC organization: (E) number of FBL-positive DFC units per nucleolus and (F) normalized DFC rim enrichment. (G–H) Three-dimensional quantification of FBL–NPM1 spatial overlap: (G) percentage of FBL-positive volume overlapping with the NPM1-positive region per nucleolus and (H) total FBL–NPM1 colocalizing volume per nucleolus. Representative three-dimensional reconstructions are shown in Videos S3–S5. A total of 163, 322, and 370 nucleoli from 99, 167, and 114 cells, respectively, and 2,028, 1,810, and 1,641 FBL-positive DFC units, respectively, were analyzed in the DMSO group and in the groups treated with SU056 at concentrations of 0.25 and 2.0 μM, respectively, across multiple experimental days. The data are presented as box-and-whisker plots overlaid with individual data points. Boxes indicate the median and interquartile range, and whiskers indicate the 1st–99th percentiles. Statistical significance was assessed using one-way analysis of variance. \**p* < 0.05, \*\**p* < 0.01, \*\*\**p* < 0.001.

SU056 treatment also altered the DFC organization. The number of FBL-positive DFC units per nucleolus decreased, accompanied by reduced DFC rim enrichment (Fig. 4E and 4F). Additional analysis showed decreases in the total FBL-positive area per nucleolus and changes in the solidity of individual FBL-positive DFC units (Figs. S5A and S5B). Three-dimensional colocalization analysis showed a decrease in the percentage of FBL-positive volume overlapping with the NPM1-positive region following SU056 treatment (Fig. 4G), together with a decrease in the total FBL–NPM1 colocalizing volume per nucleolus (Fig. 4H), as further illustrated by the reconstructions in Videos S3–S5. Together, these results show that SU056 treatment shifts multiple features of the MDA-MB-231 nucleoli, including nucleolar morphology, DFC organization, and DFC–GC spatial overlap, toward an MCF-10A-like profile.

### SU056 treatment shifts intranucleolar rRNA trafficking toward MCF-10A-like dynamics

Having found that SU056 treatment shifted the nucleolar morphology and DFC–GC organization toward an MCF-10A-like profile, we next examined whether SU056 also altered the intranucleolar dynamics of nascent rRNA. Nascent rRNA trafficking was analyzed using the same fluorescence pulse–chase approach used to compare MCF-10A and MDA-MB-231 cells, using SU056 at concentrations of 0.25 and 2.0 μM as mild and strong perturbation conditions, respectively (Fig. 5A–5C). In DMSO-treated MDA-MB-231 cells, the 5-EU signal was rapidly redistributed from the DFC-associated regions to the surrounding GC during the chase period. In contrast, SU056 treatment delayed redistribution at both concentrations, with a more pronounced delay under the strong perturbation condition (Fig. 5A–5F).

**Figure 5.**
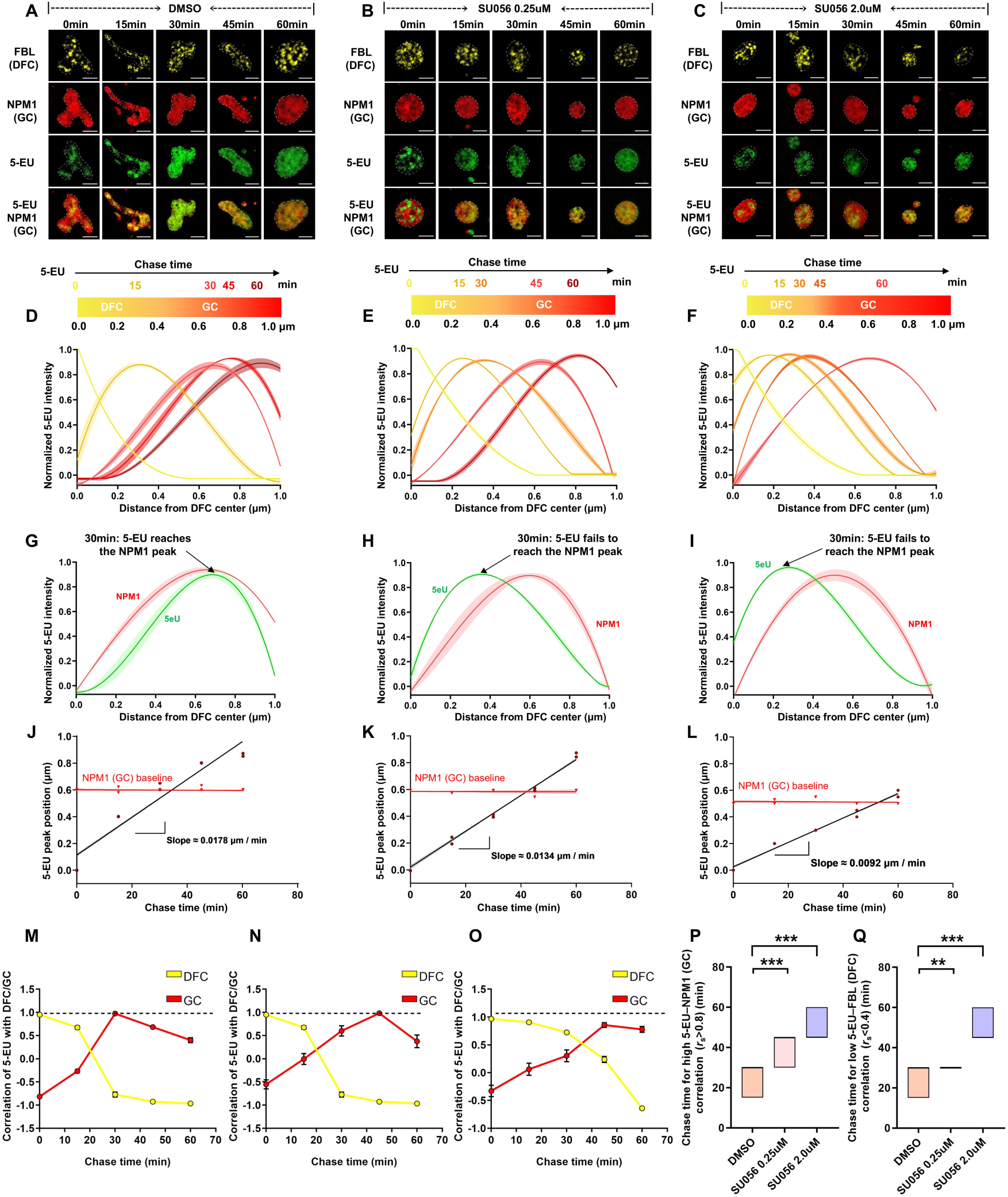
SU056 treatment shifts intranucleolar trafficking of nascent rRNA toward MCF-10A-like dynamics in MDA-MB-231 cells. (A–C) Representative fluorescence pulse–chase images showing the intranucleolar spatiotemporal distribution of nascent rRNA in MDA-MB-231 cells treated with (A) Dimethyl sulfoxide (DMSO) or SU056 at concentrations of (B) 0.25 μM and (C) 2.0 μM at the indicated chase times (0, 15, 30, 45, and 60 min). Nascent RNA was pulse-labeled with 5-ethynyl uridine (5-EU; green). Fibrillarin (FBL; yellow) and nucleophosmin 1 (NPM1; red) indicate the dense fibrillar component (DFC) and granular component (GC), respectively. Dashed lines indicate nucleolar boundaries. Scale bar = 5 μm. (D–F) Spatial profiles of normalized 5-EU fluorescence intensity in MDA-MB-231 cells treated with (D) DMSO or SU056 at concentrations of (E) 0.25 μM and (F) 2.0 μM over the 0–60-min chase period. The x-axis represents the distance from the DFC center. (G–I) Spatial profiles of normalized 5-EU (green) and NPM1 (red) fluorescence intensities at 30 min of chase in cells treated with (G) DMSO or SU056 at concentrations of (H) 0.25 μM and (I) 2.0 μM, plotted as a function of the distance from the DFC center. (J–L) Regression analysis of the displacement of the 5-EU signal peak over time in cells treated with (J) DMSO or SU056 at concentrations of (K) 0.25 μM and (L) 2.0 μM. The lines indicate linear regression fits, and the slopes represent the apparent intranucleolar translocation velocities. NPM1 peak positions are shown as a reference (red). The calculated velocities were 0.0178, 0.0134, and 0.0092 μm/min for the DMSO, 0.25 μM SU056, and 2.0 μM SU056 conditions, respectively. (M–O) Correlation analysis of 5-EU fluorescence with FBL (DFC) and NPM1 (GC) over the 0–60-min chase period in cells treated with (M) DMSO or SU056 at concentrations of (N) 0.25 μM and (O) 2.0 μM. Correlations were calculated using Spearman’s correlation coefficients. (P) Chase time required to reach a high Spearman correlation between 5-EU (nascent rRNA) and NPM1 (GC) (*r*ₛ > 0.8). (Q) Chase time required to reach a low Spearman correlation between 5-EU (nascent rRNA) and FBL (DFC) (*r*ₛ < 0.4). For (D–I), solid lines and shaded areas represent mean ± standard error of the mean. For (P–Q), data are presented as mean ± standard deviation. The numbers of nucleoli analyzed at 0, 15, 30, 45, and 60 min were 65, 79, 69, 60, and 60 for the DMSO group (38, 46, 40, 35, and 35 cells, respectively); 69, 72, 61, 70, and 60 for the 0.25 μM SU056 group (44, 46, 39, 44, and 38 cells, respectively); and 59, 66, 59, 77, and 61 for the 2.0 μM SU056 group (32, 36, 32, 42, and 33 cells, respectively). Statistical significance was assessed using a one-way analysis of variance. \*\**p* < 0.01, \*\*\**p* < 0.001.

After 30 min of the chase, the 5-EU signal in DMSO-treated cells showed substantial spatial overlap with the NPM1-positive GC region, whereas following SU056 treatment, the 5-EU signal remained more centrally distributed than the NPM1 signal (Fig. 5G–5I). Regression analysis of the displacement of the 5-EU signal peak further revealed a concentration-dependent decrease in apparent intranucleolar translocation velocity, from 0.0178 μm/min in DMSO-treated cells to 0.0134 and 0.0092 μm/min following treatment with SU056 at concentrations of 0.25 and 2.0 μM, respectively (Fig. 5J–5L).

Correlation analysis further showed that the temporal association of nascent rRNA with DFC and GC shifted following SU056 treatment (Fig. 5M–5O). In DMSO-treated cells, the 5-EU–NPM1 correlation peaked at approximately 30 min, whereas after SU056 treatment, the peak was delayed to approximately 45–60 min. Consistent with this delayed redistribution, SU056-treated cells required a longer chase time to reach a high correlation between 5-EU and NPM1 and a low correlation between 5-EU and FBL (Fig. 5P and 5Q). Together, these results demonstrate that SU056 treatment shifts the accelerated intranucleolar trafficking of nascent rRNA in MDA-MB-231 cells toward the slower dynamics observed in MCF-10A cells. This shift in rRNA dynamics occurred in parallel with SU056-induced changes in nucleolar morphology and DFC–GC organization, indicating that structural and dynamic nucleolar features can shift coordinately toward an MCF-10A-like profile.

### Transient 1,6-hexanediol treatment induces reversible cancer-like nucleolar remodeling in MCF-10A cells

The coordinated changes in nucleolar morphology, DFC–GC organization, and nascent rRNA dynamics observed in MDA-MB-231 cells suggested that these cancer-associated features might not arise independently. The nucleolus is organized as a multiphase biomolecular condensate^10,13^, in which the spatial organization of distinct nucleolar phases is closely coupled to rRNA processing and movement^18^. We therefore hypothesized that the coordinated structural and dynamic changes observed in MDA-MB-231 cells might reflect altered physicochemical organization of the nucleolus as a multiphase condensate.

To test this possibility, the molecular interactions contributing to the nucleolar condensate organization were transiently perturbed. The 1,6-hexanediol (1,6-HD) interferes with weak hydrophobic interactions associated with biomolecular condensates and has been shown to alter nucleolar organization, including the localization and spatial organization of nucleolar proteins such as NPM1 and FBL.^26,27^ Therefore, whether transient 1,6-HD treatment of non-transformed MCF-10A cells could recapitulate the cancer-associated nucleolar features observed in MDA-MB-231 cells and whether such changes were reversible after washout were investigated (Fig. 6A). Line profile analysis revealed that 1,6-HD treatment altered the spatial relationship between FBL and NPM1 signals, with increased NPM1 fluorescence at positions corresponding to FBL-positive DFC regions (Fig. 6B). Following washout, the spatial profiles of FBL and NPM1 shifted back toward those observed in the control MCF-10A cells. Treatment with 1,6-HD also induced changes in the nucleolar morphology. The individual nucleolar area increased, whereas circularity decreased, shifting both parameters toward the values observed in MDA-MB-231 cells (Fig. 6C and 6D). Additional morphological parameters, including nucleolar number and solidity, were also altered after 1,6-HD treatment (Figs. S6A and S6B). Following washout, these morphological parameters shifted back toward the control MCF-10A profile.

**Figure 6.**
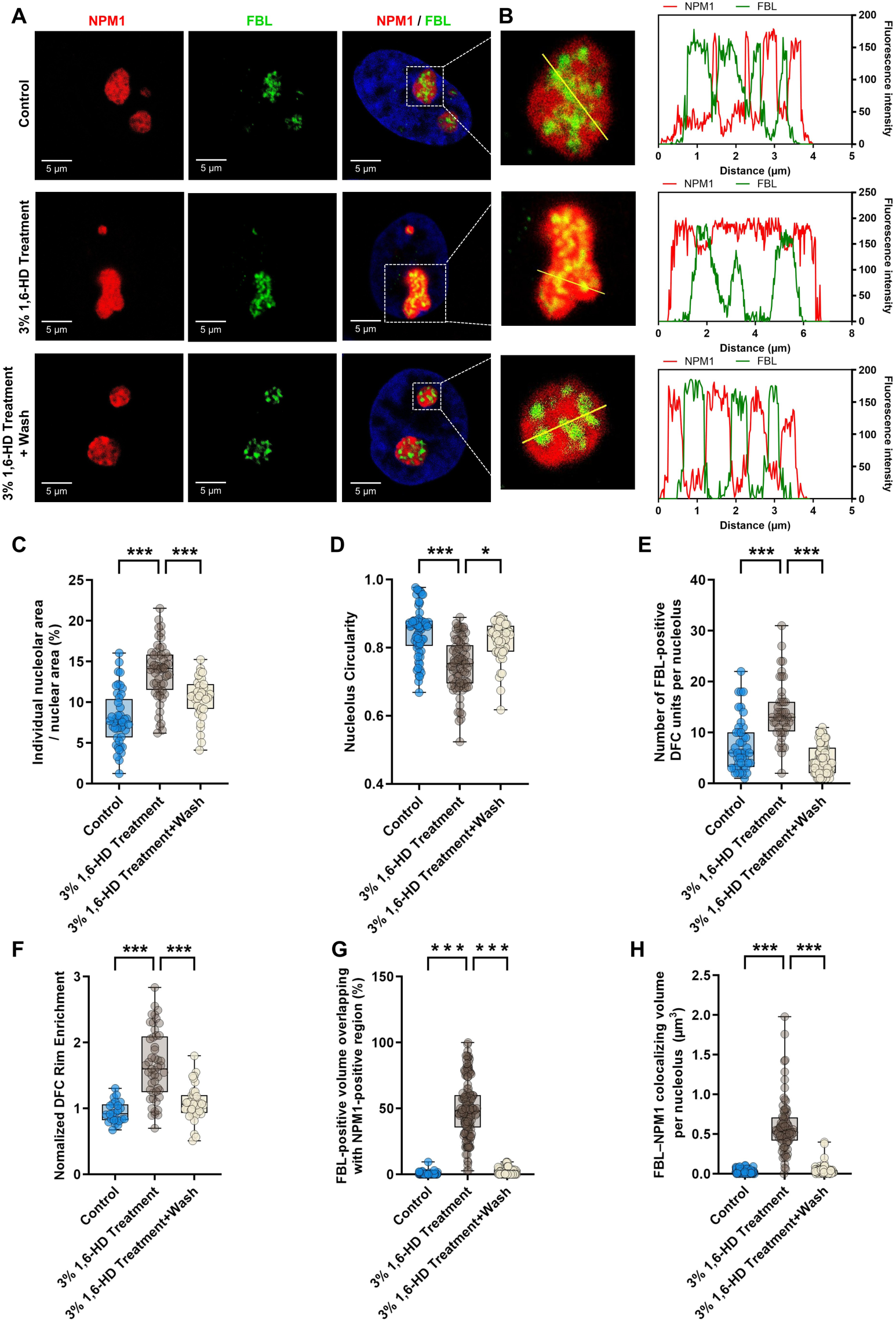
1,6-Hexanediol treatment induces reversible cancer-like nucleolar remodeling in MCF-10A cells. (A) Representative immunofluorescence images of nucleophosmin 1 (NPM1; red, granular component [GC]) and fibrillarin (FBL; green, dense fibrillar component [DFC]) in control MCF-10A cells, MCF-10A cells treated with 3% 1,6-hexanediol (1,6-HD), and cells following washout of 1,6- HD. Scale bar = 5 μm. (B) Representative line profile analysis of NPM1 (red) and FBL (green) fluorescence intensities. Yellow lines in the magnified insets corresponding to the dashed boxes in (A) indicate the scan paths for the intensity profiles shown on the right. The x-axis represents distance along the scan line (μm), and the y-axis represents fluorescence intensity. (C–D) Quantification of nucleolar morphology: (C) individual nucleolar area relative to nuclear area and (D) circularity. (E–F) Quantification of DFC organization: (E) number of FBL-positive DFC units per nucleolus and (F) normalized DFC rim enrichment. (G–H) Three-dimensional quantification of FBL–NPM1 spatial overlap: (G) percentage of FBL-positive volume overlapping with the NPM1- positive region per nucleolus and (H) total FBL–NPM1 colocalizing volume per nucleolus. Representative three-dimensional reconstructions are shown in Videos S6–S8. A total of 54, 63, and 65 nucleoli from 27, 37, and 40 cells, respectively, and 416, 945, and 1,170 FBL-positive DFC units, respectively, were analyzed in the control, 3% 1,6-HD treatment, and washout groups, respectively, across multiple experimental days. The data are presented as box- and-whisker plots overlaid with individual data points. Boxes indicate the median and interquartile range, and whiskers indicate the 1st–99th percentiles. Statistical significance was assessed using a one-way analysis of variance. \*\**p* < 0.01, \*\*\**p* < 0.001.

Consistent with the line profile analysis, quantitative analysis of DFC–GC organization showed that the number of FBL-positive DFC units per nucleolus increased following 1,6-HD treatment, together with a change in DFC rim enrichment (Fig. 6E and 6F). Additional analysis revealed changes in the total FBL-positive area per nucleolus and the solidity of individual FBL-positive DFC units (Figs. S6C and S6D). Three-dimensional colocalization analysis showed an increase in the percentage of FBL-positive volume overlapping with the NPM1-positive region and in the total FBL–NPM1 colocalizing volume per nucleolus (Fig. 6G and 6H; Videos S6–S8). These parameters also shifted back toward the control MCF-10A profile after washout.

The response of MDA-MB-231 cells to 1,6-HD was also examined. In these cells, 1,6-HD induced further changes in the nucleolar morphology and DFC–GC organization (Fig. S7). Line profile analysis revealed a marked redistribution of FBL following 1,6-HD treatment, with FBL-positive regions exhibiting a more continuous spatial distribution within the nucleolus (Figs. S7A and S7B). Consistent with this observation, quantitative analysis revealed changes in both the number of FBL-positive DFC units and the total FBL-positive area per nucleolus (Figs. S7G and S7H). In contrast, three-dimensional colocalization analysis showed no significant changes in either the percentage of FBL-positive volume overlapping with the NPM1-positive region or the total FBL–NPM1 colocalizing volume (Figs. S7K and S7L), indicating that FBL redistribution was not accompanied by a corresponding loss of NPM1 from FBL-positive regions. Following washout, changes in nucleolar morphology and FBL-positive DFC organization shifted back toward the control MDA-MB-231 profile. Thus, despite their distinct basal nucleolar profiles, both MCF-10A and MDA-MB-231 cells exhibited reversible remodeling of the nucleolar subcompartment organization in response to transient 1,6-HD perturbation. Together, these results show that transient 1,6-HD treatment induces multiple MDA-MB-231-like features of nucleolar morphology and DFC–GC organization in non-transformed MCF-10A cells and that these changes are reversible following washout. The reversible remodeling observed in both cell types further indicates that the nucleolar subcompartment organization remains responsive to physicochemical perturbations.

## DISCUSSION

In this study, cancer-associated nucleolar alterations extended beyond changes in overall morphology and involved differences in internal subcompartment organization and nascent rRNA dynamics. Compared with non-transformed MCF-10A cells, MDA-MB-231 breast cancer cells exhibited differences in nucleolar size and shape, together with altered DFC–GC organization and accelerated intranucleolar trafficking of nascent rRNA (Fig. 1–3). These features were responsive to perturbation; SU056 treatment shifted multiple structural and dynamic parameters toward the MCF-10A-like profile (Fig. 4 and 5), whereas transient 1,6-HD treatment of MCF-10A cells induced several MDA-MB-231-like nucleolar features that were reversed following washout (Fig. 6). Furthermore, in MDA-MB-231 cells, 1,6-HD altered the nucleolar morphology and DFC–GC organization, including marked changes in FBL-positive DFC organization, with these changes again shifting back toward the untreated profile following washout (Fig. S7). These findings suggest that cancer-associated nucleolar differences are not simply a collection of independent abnormalities, but reflect a broader reorganization involving both the spatial architecture and dynamic behavior of the nucleolus.

Changes in nucleolar size, number, and shape have long been recognized as characteristic features of cancer cells and have generally been discussed in relation to enhanced ribosome biogenesis and proliferative activity.^1,7,8^ Our results extend this morphological view by showing that the differences between MCF-10A and MDA-MB-231 cells also involve the spatial organization of nucleolar subcompartments. In MDA-MB-231 cells, an increase in nucleolar size and altered shape were accompanied by an increased number of FBL-positive DFC units and total FBL-positive area per nucleolus, altered DFC positioning, and greater spatial overlap between the FBL-positive DFC and NPM1-positive GC regions (Fig. 1 and 2). These observations suggest that cancer-associated changes in overall nucleolar morphology are accompanied by a broader reorganization of internal nucleolar architecture. This interpretation is consistent with the current view of the nucleolus as a multiphase condensate, in which the organization of the DFC and GC is determined by the interactions among molecularly distinct but physically coupled phases.^10,13^ Thus, the morphological differences observed in cancer nucleoli may accompany changes in the spatial organization of their internal subcompartments.

Beyond the structural organization, our results showed that cancer-associated nucleolar differences also extend to the intranucleolar dynamics of nascent rRNA. Pre-rRNA undergoes spatially ordered trafficking through nucleolar subcompartments during processing, and perturbation of this process can alter nucleolar organization.^17,18^ Consistent with a broader coupling between rRNA biogenesis and nucleolar architecture, perturbation of rRNA transcription has also been shown to alter nucleolar phase separation and structural organization.^28^ In MDA-MB-231 cells, nascent rRNA is redistributed from FBL-positive DFC regions toward NPM1-positive GC regions more rapidly than in MCF-10A cells, indicating accelerated intranucleolar trafficking (Fig. 3). The SU056 treatment slowed this accelerated redistribution, while simultaneously shifting the nucleolar morphology and DFC–GC organization toward an MCF-10A-like profile (Fig. 4 and 5). Parallel changes in structure and nascent rRNA dynamics are consistent with the emerging view that nucleolar architecture and rRNA processing and trafficking are closely interconnected.^17,18^ However, the direction of this relationship remains unresolved, as the present findings do not distinguish whether altered subcompartment organization influences rRNA trafficking, or whether changes in rRNA processing and trafficking contribute to structural organization. Therefore, our findings extend recent insights into the coupling between nucleolar architecture and rRNA dynamics to a cancer-associated context and suggest that altered nascent rRNA trafficking represents an additional dimension of nucleolar reorganization in MDA-MB-231 cells.

The parallel structural and dynamic changes observed following SU056 treatment raise the question of which molecular pathways are linked to these different aspects of nucleolar organization. YB-1 is a multifunctional RNA-binding protein involved in translational regulation and cancer progression, including breast cancer.^19,24^ Proteomic analysis of SU056-treated MDA-MB-231 cells has shown that SU056 treatment broadly affects proteins associated with protein translation and ribosomal function.^24^ Furthermore, recent evidence from another cancer model has directly linked YBX1 to ribosome biogenesis through regulation of FBL, a core component of the DFC.^29^ Thus, the nucleolar changes observed following SU056 treatment may reflect broader remodeling of molecular networks supporting ribosome biogenesis and nucleolar organization rather than alteration of a single structural component. Consistent with this hypothesis, our integrated transcriptomic and proteomic analyses identified multiple SU056-responsive factors associated with nucleolar function and rRNA processing, including DDX18, DDX21, FBL, NCL, NOP56, and NPM1 (Fig. S3). Several of these factors play established roles in different aspects of nucleolar function. For example, DDX21 coordinates rRNA transcription, processing, and modification by interacting with rRNA and small nucleolar RNAs,^30^ whereas NPM1 undergoes multivalent interactions with rRNA and nucleolar proteins, which contribute to the liquid-like organization of the GC^31^. In this context, recent work on DDX18 provides a potential connection between these molecular and structural aspects of nucleolar organization.^32^ DDX18 has been shown to contribute to nucleolar phase separation and subcompartment organization through interactions with NPM1 and nucleolar RNAs.^32^ Although the contribution of DDX18 or other individual candidates to the phenotypes observed here remains to be determined, these findings illustrate how changes in RNA-processing and ribosome-biogenesis networks could be associated with broader changes in nucleolar structure and dynamics.

The reversibility of nucleolar organization following 1,6-HD treatment provides a complementary perspective to these findings. Biomolecular condensates are dynamic assemblies that respond to changes in molecular interactions and environmental conditions.^10,13,14^ Moreover, 1,6-HD perturbs the weak hydrophobic interactions associated with biomolecular condensates and has been shown to alter nucleolar organization, including the localization and spatial organization of NPM1 and FBL.^26,27^ In MCF-10A cells, transient 1,6-HD treatment induced multiple features resembling those observed in MDA-MB-231 cells, including increased nucleolar area, reduced circularity, altered DFC organization, and increased spatial overlap between the FBL-positive DFC and NPM1-positive GC regions (Fig. 6). These changes shifted back toward the untreated MCF-10A profile following washout, indicating that these cancer-associated features can be transiently induced in non-transformed cells and are reversible following the removal of the perturbation. These changes occurred after only 5 min of 1,6-HD treatment and showed substantial recovery within 1 h after washout. This rapid timescale is consistent with the reorganization of molecular interactions underlying nucleolar subcompartmentation. The response of MDA-MB-231 cells provides an additional perspective: despite their distinct basal nucleolar organization, 1,6-HD further altered both the nucleolar morphology and DFC organization, including pronounced changes in FBL-positive DFCs, with substantial recovery toward the untreated profile following washout (Fig. S7). The rapid responsiveness and reversibility observed in both cell types are consistent with the dynamic maintenance of nucleolar subcompartmentation rather than structural fixation. Together with the multiphase nature of the nucleolus,^10,13^ these observations support the possibility that some cancer-associated nucleolar features reflect differences in perturbable physicochemical organization rather than irreversible structural alterations.

Taken together, our findings support the view that cancer-associated nucleolar alterations are an integrated reorganization that extends across morphology, internal subcompartment architecture, and nascent rRNA dynamics. Rather than representing isolated abnormalities, these features differed between MCF-10A and MDA-MB-231 cells and shifted in response to pharmacological or physicochemical perturbations. This reversibility further indicates that cancer-associated nucleolar organization retains its capacity for reorganization in response to perturbations. At the same time, the distinct nucleolar profile observed in untreated MDA-MB-231 cells suggests that this organization is maintained under basal conditions, raising the possibility that cancer-associated changes become established as a characteristic nucleolar organization across morphological, spatial, and dynamic dimensions. The molecular mechanisms that establish and maintain such an organization, including how molecular regulatory networks and physicochemical properties contribute to its formation and maintenance, remain unclear. By linking morphology, subcompartment organization, and rRNA dynamics within a common framework, our findings provide a basis for exploring how distinct nucleolar profiles are established, maintained, and remodeled across cancer types and cellular contexts.

### Limitations of the study

Although our findings provide an integrated view of cancer-associated nucleolar differences in morphology, subcompartment organization, and nascent rRNA dynamics, our analyses focused primarily on comparing MCF-10A and MDA-MB-231 cells. Therefore, whether similar nucleolar features are observed across additional breast cancer models and other cancer types remains to be determined. In addition, although changes in nucleolar morphology, DFC–GC organization, and nascent rRNA dynamics occurred in parallel, the present study does not establish causal relationships between these features. The molecular mechanisms underlying the observed nucleolar organization remain unclear. SU056 affects a broader network of ribosome-related factors, and the contributions of individual candidates identified by our integrated analysis were not directly tested, whereas 1,6-HD provided a physicochemical perturbation but did not selectively target a specific nucleolar interaction.

## RESOURCE AVAILABILITY

### Lead contact

Further information and requests for resources and reagents should be directed to Haruko Takahashi at.

### Materials availability

This study did not generate new unique reagents.

### Data and code availability

- The publicly available transcriptomic and proteomic datasets analyzed in this study are listed below. Original imaging data generated in this study are available from the lead contact upon reasonable request. The transcriptomic/gene expression datasets are accessible at the GEO under accession numbers GSE247819^20^, GSE158085^21^, GSE298177^22^, and GSE271749^23^. The mass spectrometry proteomics data are available in the PRIDE Archive (http://www.ebi.ac.uk/pride/archive/) under the dataset identifier PXD042380.
- This article does not report the original code.
- Any additional information required to reanalyze the data reported in this article is available from the lead contact upon request.

## ACKNOWLEDGEMENTS

This study was partly supported by the Hiroshima University Grant-in-Aid for Exploratory Research (to K.-D.F.), JSPS KAKENHI Grant Number 24K157470A and a Grant-in-Aid from the Graduate School of Integrated Sciences for Life, Hiroshima University (to H.T.).

## AUTHOR CONTRIBUTIONS

Conceptualization, K.-D.F., Y.K., and H.T.; methodology, K.-D.F., Y.K., and H.T.; investigation, K.-D.F., W.-H.W., G.K., Y.-D.-L.L. and H.T.; writing—original draft, K.-D.F.; writing—review & editing, Y.K. and H.T.; supervision, Y.K. and H.T. All authors interpreted the results and read and approved the final version of the manuscript.

## DECLARATION OF GENERATIVE AI AND AI-ASSISTED TECHNOLOGIES IN THE WRITING PROCESS

No generative AI or AI-assisted technologies were used in the writing or preparation of this manuscript.

## SUPPLEMENTAL INFORMATION

Document S1. Figures S1–S7

Videos S1–S8. Imaris (version 9.9.0; Oxford Instruments) 3D reconstructions showing the spatial overlap of NPM1 and FBL. Videos S1 and S2 are related to Figure 2; Videos S3–S5 are related to Figure 4; and Videos S6–S8 are related to Figure 6.

## STAR★METHODS

### KEY RESOURCES TABLE

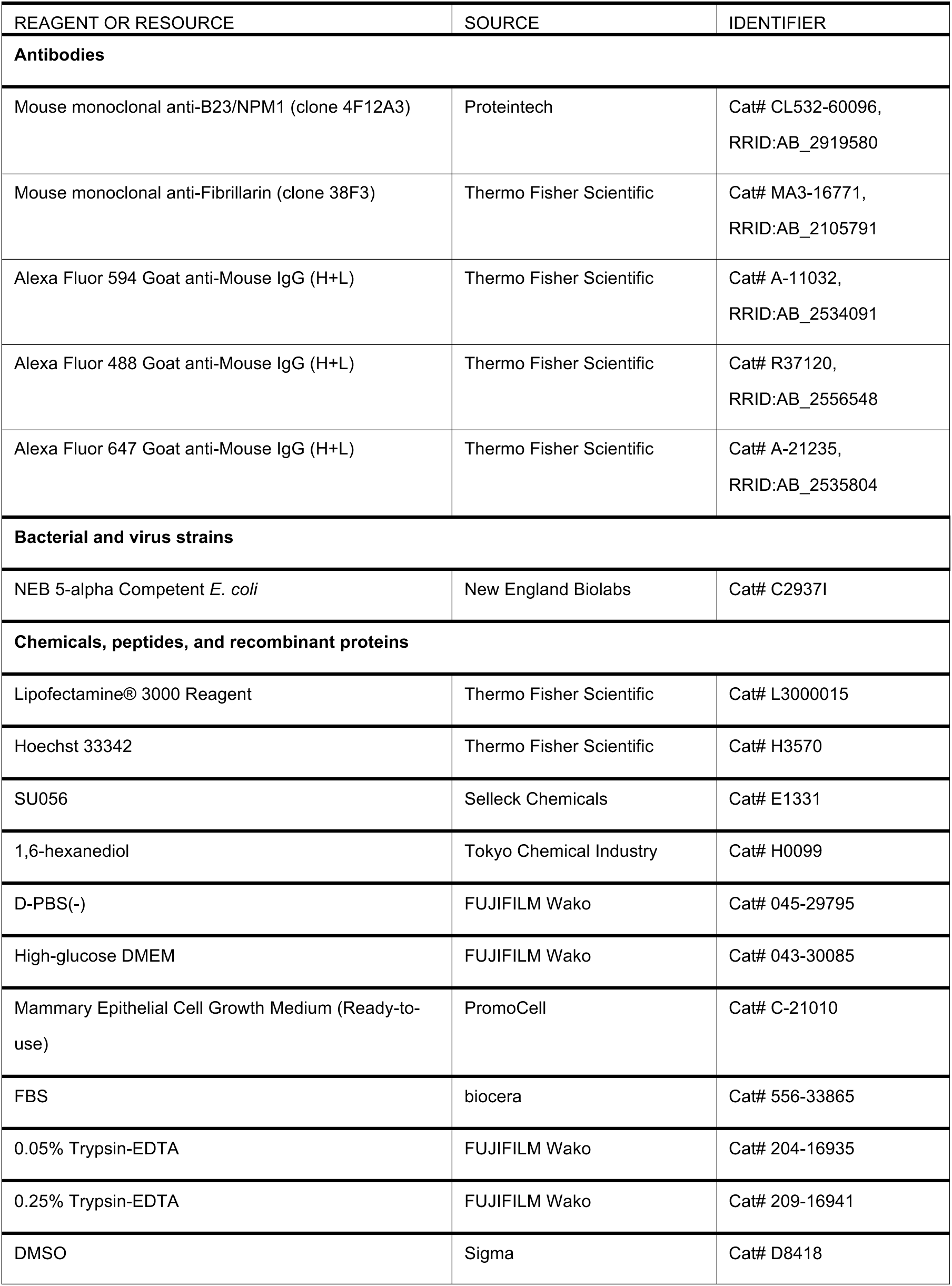
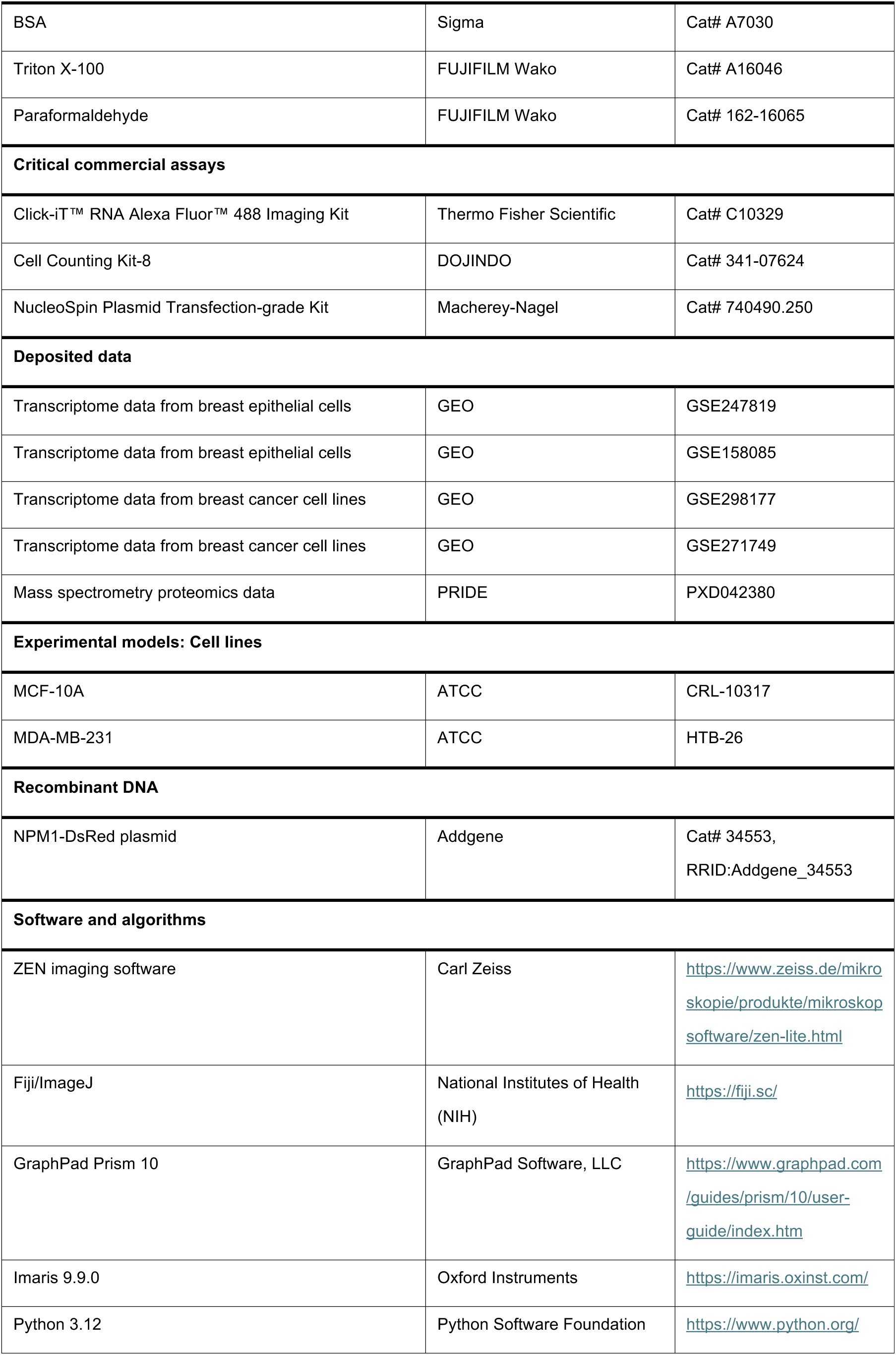
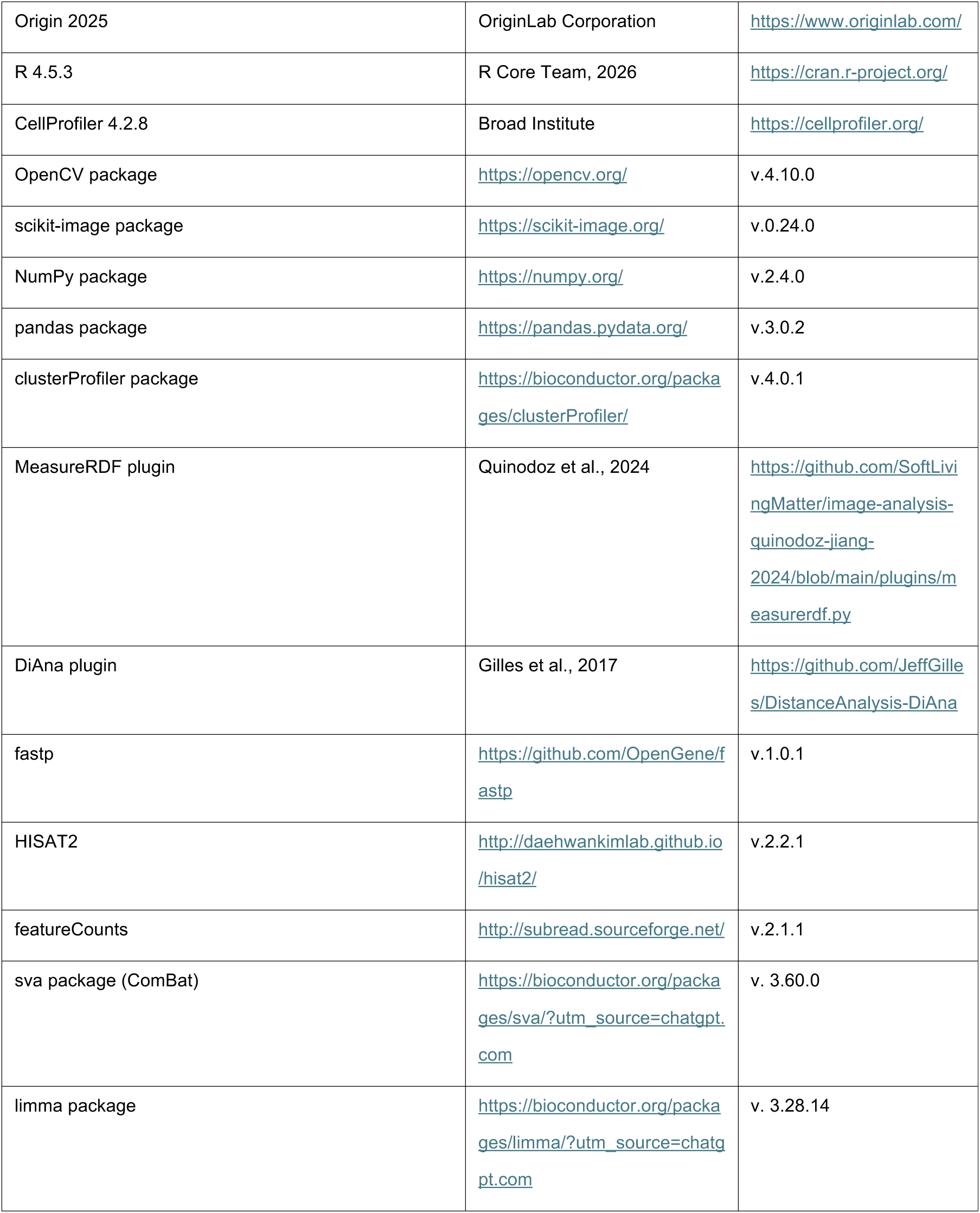

| REAGENT OR RESOURCE | SOURCE | IDENTIFIER |
| --- | --- | --- |
| **Antibodies** |  |  |
| Mouse monoclonal anti-B23/NPM1 (clone 4F12A3) | Proteintech | Cat# CL532-60096,<br>RRID:AB_2919580 |
| Mouse monoclonal anti-Fibrillarin (clone 38F3) | Thermo Fisher Scientific | Cat# MA3-16771,<br>RRID:AB_2105791 |
| Alexa Fluor 594 Goat anti-Mouse IgG (H+L) | Thermo Fisher Scientific | Cat# A-11032,<br>RRID:AB_2534091 |
| Alexa Fluor 488 Goat anti-Mouse IgG (H+L) | Thermo Fisher Scientific | Cat# R37120,<br>RRID:AB_2556548 |
| Alexa Fluor 647 Goat anti-Mouse IgG (H+L) | Thermo Fisher Scientific | Cat# A-21235,<br>RRID:AB_2535804 |
| **Bacterial and virus strains** |  |  |
| NEB 5-alpha Competent *E. coli* | New England Biolabs | Cat# C2937I |
| **Chemicals, peptides, and recombinant proteins** |  |  |
| Lipofectamine® 3000 Reagent | Thermo Fisher Scientific | Cat# L3000015 |
| Hoechst 33342 | Thermo Fisher Scientific | Cat# H3570 |
| SU056 | Selleck Chemicals | Cat# E1331 |
| 1,6-hexanediol | Tokyo Chemical Industry | Cat# H0099 |
| D-PBS(-) | FUJIFILM Wako | Cat# 045-29795 |
| High-glucose DMEM | FUJIFILM Wako | Cat# 043-30085 |
| Mammary Epithelial Cell Growth Medium (Ready-to-use) | PromoCell | Cat# C-21010 |
| FBS | biocera | Cat# 556-33865 |
| 0.05% Trypsin-EDTA | FUJIFILM Wako | Cat# 204-16935 |
| 0.25% Trypsin-EDTA | FUJIFILM Wako | Cat# 209-16941 |
| DMSO | Sigma | Cat# D8418 |
| BSA | Sigma | Cat# A7030 |
| Triton X-100 | FUJIFILM Wako | Cat# A16046 |
| Paraformaldehyde | FUJIFILM Wako | Cat# 162-16065 |
| **Critical commercial assays** |  |  |
| Click-iT™ RNA Alexa Fluor™ 488 Imaging Kit | Thermo Fisher Scientific | Cat# C10329 |
| Cell Counting Kit-8 | DOJINDO | Cat# 341-07624 |
| NucleoSpin Plasmid Transfection-grade Kit | Macherey-Nagel | Cat# 740490.250 |
| **Deposited data** |  |  |
| Transcriptome data from breast epithelial cells | GEO | GSE247819 |
| Transcriptome data from breast epithelial cells | GEO | GSE158085 |
| Transcriptome data from breast cancer cell lines | GEO | GSE298177 |
| Transcriptome data from breast cancer cell lines | GEO | GSE271749 |
| Mass spectrometry proteomics data | PRIDE | PXD042380 |
| **Experimental models: Cell lines** |  |  |
| MCF-10A | ATCC | CRL-10317 |
| MDA-MB-231 | ATCC | HTB-26 |
| **Recombinant DNA** |  |  |
| NPM1-DsRed plasmid | Addgene | Cat# 34553,<br>RRID:Addgene_34553 |
| **Software and algorithms** |  |  |
| ZEN imaging software | Carl Zeiss | https://www.zeiss.de/mikroskopie/produkte/mikroskopsoftware/zen-lite.html |
| Fiji/ImageJ | National Institutes of Health (NIH) | https://fiji.sc/ |
| GraphPad Prism 10 | GraphPad Software, LLC | https://www.graphpad.com/guides/prism/10/user-guide/index.htm |
| Imaris 9.9.0 | Oxford Instruments | https://imaris.oxinst.com/ |
| Python 3.12 | Python Software Foundation | https://www.python.org/ |
| Origin 2025 | OriginLab Corporation | https://www.originlab.com/ |
| R 4.5.3 | R Core Team, 2026 | https://cran.r-project.org/ |
| CellProfiler 4.2.8 | Broad Institute | https://cellprofiler.org/ |
| OpenCV package | https://opencv.org/ | v.4.10.0 |
| scikit-image package | https://scikit-image.org/ | v.0.24.0 |
| NumPy package | https://numpy.org/ | v.2.4.0 |
| pandas package | https://pandas.pydata.org/ | v.3.0.2 |
| clusterProfiler package | https://bioconductor.org/packages/clusterProfiler/ | v.4.0.1 |
| MeasureRDF plugin | Quinodoz et al., 2024 | https://github.com/SoftLivingMatter/image-analysis-quinodoz-jiang-2024/blob/main/plugins/measurerdf.py |
| DiAna plugin | Gilles et al., 2017 | https://github.com/JeffGilles/DistanceAnalysis-DiAna |
| fastp | https://github.com/OpenGene/fastp | v.1.0.1 |
| HISAT2 | http://daehwankimlab.github.io/hisat2/ | v.2.2.1 |
| featureCounts | http://subread.sourceforge.net/ | v.2.1.1 |
| sva package (ComBat) | https://bioconductor.org/packages/sva/?utm_source=chatgpt.com | v. 3.60.0 |
| limma package | https://bioconductor.org/packages/limma/?utm_source=chatgpt.com | v. 3.28.14 |

### EXPERIMENTAL MODEL AND STUDY PARTICIPANT DETAILS

#### Cell lines

Human breast adenocarcinoma MDA-MB-231 cells were obtained from the American Type Culture Collection (ATCC) and maintained in D-MEM supplemented with 10% FBS. Human breast epithelial cell line MCF-10A cells were obtained from the ATCC and maintained in mammary epithelial cell growth medium (PromoCell). All cell cultures in this study were maintained without the use of antibiotics or antimycotics. Cells were harvested after 3 min of incubation with 0.25% trypsin-EDTA (for MDA-MB-231) or 0.05% trypsin-EDTA (for MCF-10A), collected by centrifugation (400 × g, 3 min) prior to resuspension in the medium, and cultured at 37°C with 5% CO_2_. MCF-10A and MDA-MB-231 cells were used for the experiments after 10–15 passages of routine subculture.

### METHOD DETAILS

#### Plasmid preparation

The NPM1-DsRed plasmids were obtained from Addgene. For amplification, the plasmids were transformed into NEB 5-alpha competent Escherichia coli (New England Biolabs) and cultured overnight. Subsequently, the plasmids were extracted and purified using the NucleoSpin Plasmid Transfection-grade kit (Macherey-Nagel) according to the manufacturer’s instructions.

#### Transient transfection

The NPM1-DsRed plasmid was used to express fluorescently labeled NPM1. The plasmids were transfected into MCF-10A and MDA-MB-231 cells using Lipofectamine™ 3000 (Thermo Fisher Scientific). Cells were seeded into 35 mm glass-bottom dishes (Matsunami Glass, Cat# D11130H). For experiments involving immunofluorescence staining, cells were fixed at 48 h post-transfection. For experiments involving nascent rRNA imaging, cells were transfected with the NPM1-DsRed plasmid and subsequently subjected to 5-EU labeling and immunofluorescence staining as described below.

#### Immunohistochemistry

For immunofluorescence staining, cells in glass bottom dishes (35 mm) were fixed with 4% paraformaldehyde (PFA, FUJIFILM Wako) for 15 min at room temperature. After washing PBS three times, the cells were permeabilized with 0.5% Triton X-100 (FUJIFILM Wako) in PBS for 15 min at room temperature, and then blocked in 1% bovine serum albumin (BSA, Sigma) in PBS overnight at 4°C. For NPM1 immunostaining, non-transfected cells were incubated overnight at 4°C with a mouse monoclonal anti-NPM1 primary antibody (Proteintech, 1:1000) diluted in 1% BSA solution. After washing with 1% BSA solution, cells were incubated with Alexa Fluor 594 Goat anti-Mouse IgG (H+L) (Thermo Fisher Scientific, 1:500) for 3 h at room temperature in the dark.

For NPM1–FBL colocalization analysis, NPM1-DsRed-expressing cells were incubated overnight at 4°C with a mouse monoclonal anti-Fibrillarin (FBL) primary antibody (Thermo Fisher Scientific, 1:500) diluted in 1% BSA solution. After washing with 1% BSA solution, cells were incubated with Alexa Fluor 488 Goat anti-Mouse IgG (H+L) (Thermo Fisher Scientific, 1:500) for 3 h at room temperature in the dark. NPM1 was visualized through the intrinsic fluorescence of the NPM1-DsRed fusion protein.

Following secondary antibody incubation, cells were washed once with PBS and counterstained with Hoechst 33342 (Thermo Fisher Scientific, 1:1000) for 10 min under light shielding, followed by three washes with PBS for 10 min each with agitation. Images were acquired using a laser scanning confocal microscope (Carl Zeiss, LSM700).

#### 5-EU imaging of nascent rRNA synthesis in the nucleolus

To visualize nascent rRNA synthesis, the 5-ethynyl uridine (5-EU) labeling method was performed using the Click-iT™ RNA Alexa Fluor™ 488 Imaging Kit (Thermo Fisher Scientific). MCF-10A and MDA-MB-231 cells were seeded on 35 mm glass-bottom dishes at 50% confluency one day prior to the experiment. Reagent volumes were maintained at 500 μL per dish. A 1 mM solution of 5-EU was prepared in cell culture medium and added to the cells. Cells were incubated for 30 min at 37°C with 5% CO_2_ (referred to as the “pulse”). Following the pulse, the medium containing 5-EU was removed, and cells were rapidly washed twice with PBS to remove residual 5-EU. Fresh, pre-warmed culture medium was then added, and cells were incubated for various chase periods (0, 15, 30, 45, and 60 min) at 37°C with 5% CO_2_. All solutions used in the pulse and chase steps were maintained at 37°C using a heat block to minimize temperature-induced effects on RNA transcription and processing.

For fixation, cells were treated with 4% paraformaldehyde (PFA, FUJIFILM Wako) in PBS for 15 min at room temperature. Fixed cells were washed three times with PBS and permeabilized with 0.5% Triton X-100 (FUJIFILM Wako) in PBS for 15 min. To combine immunofluorescence with 5-EU imaging, NPM1-DsRed-expressing cells were immunostained for FBL using a mouse monoclonal anti-Fibrillarin (FBL) primary antibody (Thermo Fisher Scientific, 1:500) and Alexa Fluor 647 Goat anti-Mouse IgG (H+L) secondary antibody (Thermo Fisher Scientific, 1:500), as described above, prior to the click chemistry reaction. The click chemistry reaction was performed according to the manufacturer’s instructions. Cells were incubated with the reaction cocktail containing Alexa Fluor™ 488 azide for 30 min. Subsequently, cells were washed once with the reaction rinse buffer provided in the kit and once with PBS before imaging.

#### SU056 treatment

Stock solutions of SU056 (Selleck Chemicals) were prepared by dissolving the compound in dimethyl sulfoxide (DMSO, Sigma). To evaluate the effects of SU056 across a range of concentrations, MDA-MB-231 cells were initially treated with SU056 at a series of final concentrations (0.1, 0.25, 0.5, 1.0, and 2.0 μM) diluted in fresh growth medium for 48 h at 37°C. Then, cell viability was assessed using the Cell Counting Kit-8 (DOJINDO) according to the manufacturer’s instructions. After addition of the CCK-8 reagent, cells were incubated for 2 h at 37°C, and absorbance was measured at 450 nm. Cell viability was expressed relative to the DMSO-treated vehicle control, which was set to 100%. Based on cell viability and nucleolar morphology analyses, SU056 concentrations of 0.25 and 2.0 μM were selected for subsequent experiments to represent mild and strong perturbation conditions, respectively. For the negative control group, cells were treated with an equivalent volume of DMSO (vehicle control) to account for potential solvent effects.

#### 1,6-hexanediol treatment

Stock solutions of 1,6-hexanediol (Tokyo Chemical Industry) were prepared by dissolving the compound in pure water. MCF-10A and MDA-MB-231 cells were treated with 3% 1,6-hexanediol (final concentration) in fresh growth medium for 5 min at 37°C. After treatment, the dishes were washed with fresh medium three times and then incubated for 1 h at 37°C. For the negative control group, cells were treated with an equivalent volume of pure water (vehicle control) to account for potential solvent effects.

#### Image segmentation and morphology data processing

To evaluate the morphology of nucleolar sub-compartments, a systematic analysis was performed on hundreds of multi-channel immunofluorescence images using Python, leveraging OpenCV, Scikit-image, and NumPy. This approach allowed for the consistent and efficient quantification of the Granular Component (GC, labeled by NPM1) and the Dense Fibrillar Component (DFC, labeled by Fibrillarin) across the entire dataset.

The analysis focused on two primary nucleolar substructures: the Granular Component (GC, NPM1) and the Dense Fibrillar Component (DFC, Fibrillarin). For each image in the dataset, the NPM1 channel was first converted to grayscale and segmented using Otsu’s automatic thresholding to generate binary masks. Individual nucleoli were then identified and isolated through automated contour extraction algorithms, with size-based filtering applied to exclude noise and small artifacts. For each segmented nucleolus, a comprehensive set of GC morphological features was quantified, including count, area, estimated volume, eccentricity, circularity, and solidity.

Subsequently, the green channel corresponding to Fibrillarin within each nucleolus was extracted and locally thresholded to quantify DFC features, including spot count, area, edge enrichment, estimated volume, eccentricity, and solidity. DFC rim enrichment was calculated by dividing the fraction of total Fibrillarin intensity localized within the nucleolar rim by the area fraction of the rim relative to the entire GC object. Investigation of a range of rim widths found that the outer 20% of the GC rim provided the best discrimination between MDA-MB-231 and MCF-10A cells.

Estimated Volume (*V_N_*): Calculated assuming a spherical geometry based on the projected area (*A*) using the formula: 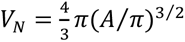

Eccentricity (*e*): Derived from a fitted ellipse as 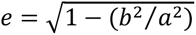, where *a* and *b* are the semi-major and semi-minor axes.

Solidity: Defined as the ratio of the object’s contour area to its convex hull area *Solidity* = *Area_contour_⁄Area_convex ℎull_*

Circularity: To measure the degree of roundness and boundary smoothness, circularity was calculated using the formula *Circularity* = 4*π*(*A*⁄*P*^2^), here *P* represents the perimeter.

DFC rim enrichment (*E_rim_*): Measures the spatial distribution bias of FBL units relative to the nucleolar boundary. It was calculated as the formula: 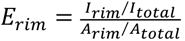, where *I_rim_* and *I_total_* represent the integrated fibrillarin intensity within the rim and the entire nucleolus, respectively.

The *A_rim_* and *A_total_* represent the area of the rim and the entire nucleolus, Quantitative measurements obtained from individual nucleoli were compiled into structured Pandas DataFrames for downstream analysis. These datasets were then subjected to statistical comparisons across experimental groups to evaluate differences in nucleolar morphological parameters.

#### Three-dimensional reconstruction

Confocal z-stacks, each comprising 15 optical sections, were acquired using an LSM 700 confocal microscope (Carl Zeiss) and imported into Imaris. Three-dimensional surface reconstructions of NPM1-positive nucleolar regions and FBL-positive dense fibrillar component (DFC) structures were generated using the Surfaces function. NPM1 and FBL structures were displayed in red and green, respectively. Colocalization between the NPM1 and FBL channels was analyzed using the Coloc module, and the identified colocalizing regions were displayed in yellow. Movies showing the reconstructed structures from different viewing angles were generated in Imaris and provided as supplementary videos.

#### Line-profile analysis of FBL and NPM1 fluorescence

Line-profile analysis of FBL and NPM1 fluorescence was performed using Fiji/ImageJ. For each representative nucleolus, a straight-line region of interest (ROI) was manually drawn across the nucleolus, spanning the nucleolar region from one side to the other. A fixed line width of 3 pixels was used for all measurements. The same line ROI was applied to the FBL and NPM1 fluorescence channels to ensure that the intensity profiles were obtained along an identical spatial path. Fluorescence intensity values were extracted separately from the FBL (green) and NPM1 (red) channels using the Plot Profile function in Fiji/ImageJ. The extracted fluorescence intensities were plotted as a function of physical distance along the scan path, with pixel positions converted to micrometers according to the spatial calibration of the images. Fluorescence intensity values were presented without normalization, and the distance along the scan path was not normalized. The yellow lines in the magnified images indicate the scan paths used to generate the corresponding fluorescence intensity profiles.

#### Three-dimensional subnucleolar co-localization analysis

To evaluate the spatial relationship between nucleolar subcompartments, the DiAna plugin ^33^ for Fiji/ImageJ was used to perform object-based three-dimensional colocalization analysis of the DFC (FBL) and GC (NPM1) compartments. Original CZI confocal z-stacks were analyzed as complete 3D stacks. The FBL and NPM1 fluorescence channels were analyzed from 3D image stacks, and FBL- and NPM1-positive regions were segmented into individual 3D objects using the segmentation procedures implemented in DiAna. The segmented FBL and NPM1 objects were then compared in three-dimensional space to identify their spatially overlapping regions and calculate the corresponding co-localizing volumes.

For each individual nucleolus, two parameters were quantified: (1) the total co-localizing volume between the FBL and NPM1 compartments and (2) the percentage of FBL volume overlapping with the NPM1-positive region. The co-localizing volume was determined from the three-dimensional spatial intersection of the segmented FBL and NPM1 objects.

Percentage of FBL volume overlapping with NPM1 (*P*_overlap_): Quantified as the proportion of the FBL compartment shared with NPM1 using the formula: *P*_overlap_ = (*V_FBL_*_∩*NPM*1_/*V_FBL_*_,*total*_) × 100%. Here, *V_FBL_*_∩*NPM*1_ represents the 3D volume shared by the FBL- and NPM1-positive objects, and *V_FBL_*_,*total*_ is the entire 3D volume of the FBL-positive compartment within the individual nucleolus.

FBL–NPM1 co-localizing volume (*V_FBL_*_∩*NPM*1_): Volume of FBL∩NPM1. These measurements were performed for individual nucleoli and used to quantify the spatial overlap between FBL-positive DFC and NPM1-positive GC regions across experimental groups.

#### Radial distribution function (RDF) estimation of 5-EU labeled rRNA

To spatially resolve the site of nascent rRNA synthesis relative to the nucleolar architecture, Radial Distribution Function (RDF) analysis was performed using the “MeasureRDF” plugin developed for CellProfiler ^18^. This method quantitatively characterizes the average radial intensity profile of the 5-EU signal as a function of distance from the DFC.

For this analysis, the centroids of individual DFC units (Fibrillarin-positive puncta) were defined as the reference point sources (r=0). This computational method provides a robust, averaged estimate of the 5-EU signal distribution relative to the DFC center while accounting for the spatial contribution of neighboring units. Radial distances were converted to micrometers using the spatial calibration of the images. The resulting radial intensity profiles were min-max normalized to visualize and compare the spatial dynamics of rRNA synthesis across experimental groups. The 5-EU peak position was identified directly from each MeasureRDF output profile as the radial coordinate corresponding to the maximum intensity, representing the distance from the DFC center to the location of maximal 5-EU enrichment.

#### Quantification of nascent rRNA trafficking kinetics

Apparent intranucleolar translocation of 5-EU-labeled nascent rRNA was assessed by tracking radial peak positions across chase times. Peak positions were obtained from the RDF profiles as described above. Radial displacement between chase times *t*_1_ and *t*_2_ was defined as:

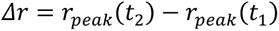

For each experimental group, peak position was regressed against chase time using simple linear regression in GraphPad Prism (v10):

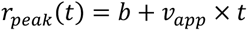

Here, *r_peak_*(*t*) denotes the radial distance of the 5-EU peak from the DFC center at time *t*, and *b* is the intercept. The regression slope, *v_app_*, was defined as the apparent intranucleolar translocation velocity. Distances were expressed in μm and chase times in min, yielding *v_app_* in μm/min.

#### Correlation analysis of nascent rRNA with DFC and GC markers

Correlations between the radial distributions of 5-EU and FBL or NPM1 were quantified using Spearman’s rank correlation in GraphPad Prism. Fluorescence intensities at matched radial distances in the RDF profiles were paired to calculate Spearman’s correlation coefficient (*r*ₛ) for 5-EU–FBL and 5-EU–NPM1 at each chase time (0, 15, 30 and 45 min for Fig. 3; 0, 15, 30, 45 and 60 min for Fig. 5).

High 5-EU–NPM1 correlation and low 5-EU–FBL correlation were defined operationally as *r*ₛ > 0.8 and *r*ₛ < 0.4, respectively. The time to reach each threshold was defined as the earliest measured chase time at which the correlation coefficient met the corresponding criterion. If the criterion was not met within the corresponding observation period (up to 45 min for Fig. 3 and 60 min for Fig. 5), the threshold was recorded as not reached within the observation period.

**The sample sizes analyzed for each experiment were as follows:**

- **Nucleolar GC morphology (NPM1):** Baseline Characterization (Fig. 1): Quantitative analysis was performed on 492 cells (*n* = 950 nucleoli) for MCF-10A, 500 cells (*n* = 821 nucleoli) for MDA-MB-231. SU056 Intervention (Fig. 4): For the evaluation of SU056 treatment on nucleolar morphology in MDA-MB-231 cells, the dataset included: DMSO vehicle control (99 cells, *n* = 163 nucleoli), 0.25 μM SU056 (167 cells, *n* = 322 nucleoli), and 2.0 μM SU056 (114 cells, *n* = 370 nucleoli). 1,6-Hexanediol Intervention (Fig. 6): For the evaluation of 1,6-HD treatment on nucleolar morphology in MCF-10A cells, the dataset included: control group (27 cells, *n* = 54 nucleoli), 3% 1,6-HD treatment group (37 cells, *n* = 63 nucleoli), and 3% 1,6-HD treatment + wash group (40 cells, *n* = 65 nucleoli). Dose-Response Screening (Fig. S4): For dose-dependent morphological profiling, the following sample sizes were analyzed: DMSO control (137 cells, *n* =234 nucleoli); 0.1 μM SU056 (100 cells, *n* =136 nucleoli); 0.25 μM SU056 (141 cells, *n* =223 nucleoli); 0.5 μM SU056 (100 cells, n =147 nucleoli); 1.0 μM SU056 (111 cells, *n* =200 nucleoli); and 2.0 μM SU056 (101 cells, *n* =191 nucleoli). 1,6-Hexanediol Intervention (Fig. S7): To evaluate the effects of 1,6-HD treatment on nucleolar morphology in MDA-MB-231 cells, the dataset included: control group (37 cells, *n* = 60 nucleoli), 3% 1,6-HD treatment group (42 cells, *n* = 67 nucleoli), and 3% 1,6-HD treatment + wash group (43 cells, *n* = 70 nucleoli).
- **Sub-nucleolar DFC architecture (FBL):** Baseline Characterization (Fig. 2, Fig. S2): Analysis included 52 MCF-10A cells (100 nucleoli; *N_FBL_* = 770 DFCs) and 54 MDA-MB-231 cells (89 nucleoli; *N_FBL_* = 1,641 DFCs). SU056 Intervention (Fig. 4, Fig. S5): To quantify the restoration of DFC organization, analysis included 99 cells (163 nucleoli; *N_FBL_* =2028 DFCs) for the DMSO control, 167 cells (322 nucleoli; *N_FBL_* =1810 DFCs) for the 0.25 μM SU056 group, and 114 cells (370 nucleoli; *N_FBL_* =1641 DFCs) for the 2.0 μM SU056 group. 1,6-Hexanediol Intervention (Fig. 6, Fig. S6): To evaluate the effects of 1,6-HD treatment in MCF-10A cells, analysis included 27 cells (54 nucleoli; *N_FBL_* = 416 DFCs) for the control group, 37 cells (63 nucleoli; *N_FBL_* = 945 DFCs) for the 3% 1,6-HD treatment group, and 40 cells (65 nucleoli; *N_FBL_* = 1170 DFCs) for the 3% 1,6-HD treatment + wash group. 1,6-Hexanediol Intervention (Fig. S7): To assess the effects of 1,6-HD treatment on DFC organization in MDA-MB-231 cells, analysis included 37 cells (60 nucleoli; *N_FBL_* = 1110 DFCs) for the control group, 42 cells (67 nucleoli; *N_FBL_* = 1246 DFCs) for the 3% 1,6-HD treatment group, and 43 cells (70 nucleoli; *N_FBL_* = 1330 DFCs) for the 3% 1,6-HD treatment + wash group.
- **Spatiotemporal rRNA trafficking kinetics (Figures 3 & 5):** Quantification of nascent rRNA dynamics was performed on individual nucleoli at sequential pulse-chase timepoints. For baseline kinetics (0-45 min chase; Fig. 3), the sample sizes at 0, 15, 30, and 45 min were: MCF-10A (*n* = 60, 59, 67, and 60 nucleoli from 31, 31, 35, and 31 cells, respectively); MDA-MB-231 (*n* = 79, 85, 87, and 72 nucleoli from 48, 52, 53, and 44 cells, respectively). For SU056 intervention (0-60 min chase; Fig. 5), MDA-MB-231 cells were quantified at 0, 15, 30, 45, and 60 min with the following sample sizes: DMSO control, 38, 46, 40, 35, and 35 cells (*n* = 65, 79, 69, 60, and 60 nucleoli, respectively); 0.25 μM SU056, 44, 46, 39, 44, and 38 cells (*n* = 69, 72, 61, 70, and 60 nucleoli, respectively); 2.0 μM SU056, 32, 36, 32, 42, and 33 cells (*n* = 59, 66, 59, 77, and 61 nucleoli, respectively).

#### Integrated multi-omics analysis of ribosome biogenesis

**RNA-seq data acquisition and preprocessing:** Raw RNA-sequencing data for normal mammary epithelial cells (MCF 10A) and breast cancer cell lines (MDA-MB-231) were retrieved from the Gene Expression Omnibus (GEO) database under accession numbers GSE247819^20^ and GSE158085^21^ (MCF 10A), and GSE298177^22^ and GSE271749^23^ (MDA-MB-231). To ensure data robustness, all datasets were processed using a standardized preprocessing workflow. Briefly, raw sequencing reads were subjected to quality control and adapter trimming using Fastp, followed by alignment to the human reference genome GRCh38 with HISAT2. Gene-level expression was quantified using FeatureCounts, converted to Transcripts Per Million (TPM) for normalization, and subsequently corrected for non-biological technical variation using the ComBat function implemented in the sva R package to remove batch effects.

**Proteomic data acquisition:** Mass spectrometry (MS)-based proteomic data from SU056-treated MDA-MB-231 cells were obtained from the supplementary dataset of Dheeraj *et al.* (2024)^24^ and integrated with transcriptomic data for comparative analysis.

To systematically evaluate nucleolar function, we utilized the 331 ribosome biogenesis–related genes (RBGS) defined by Zang et al. (2024)^25^. Differential expression analysis was performed using limma, with significance defined as an absolute log2-fold change (∣log_2_FC∣) >1 and *P* < 0.05. Principal Component Analysis (PCA) was employed to verify sample clustering and ensure experimental consistency. To interpret the biological implications of the identified DEGs, Gene Ontology (GO) functional enrichment analysis was conducted using the clusterProfiler package, categorizing results into Biological Process (BP), Cellular Component (CC), and Molecular Function (MF).

### QUANTIFICATION AND STATISTICAL ANALYSIS

Statistical analyses were performed using GraphPad Prism. Data are presented as mean ± standard deviation or standard error of the mean, as specified in the figure legends. Two independent groups were compared using an unpaired, two-tailed Student’s *t*-test, whereas comparisons among three or more groups were performed using analysis of variance (ANOVA). Spearman’s rank correlation and simple linear regression were performed as described above. In all cases, a *P*-value of less than 0.05 was considered statistically significant.

