## Supplementary Information for "Cancer-associated nucleolar morphological changes coincide with altered subcompartment organization and accelerated nascent rRNA trafficking in breast cancer cells"

### Index

**Supplemental figures**

Figure S1. Relationship between nucleolar area and the number of FBL-positive DFC units.

Figure S2. Differences in the size and shape of individual FBL-positive DFC units in MCF-10A and MDA-MB-231 cells.

Figure S3. Integrated transcriptomic and proteomic analysis identifies SU056-responsive ribosome-related factors in MDA-MB-231 cells.

Figure S4. Effects of increasing SU056 concentrations on cell viability and nucleolar morphology in MDA-MB-231 cells.

Figure S5. Additional quantitative assessment of DFC organization following SU056 treatment in MDA-MB-231 cells.

Figure S6. Additional quantitative assessment of nucleolar morphology and DFC organization following 1,6-hexanediol treatment in MCF-10A cells.

Figure S7. Effects of 1,6-hexanediol treatment on nucleolar morphology and DFC–GC organization in MDA-MB-231 cells.

**Supplemental videos**

Video S1. Three-dimensional visualization of DFC and GC organization in an MCF-10A cell.

Video S2. Three-dimensional visualization of DFC and GC organization in an MDA-MB-231 cell.

Video S3. Three-dimensional visualization of DFC and GC organization in a DMSO-treated MDA-MB-231 cell.

Video S4. Three-dimensional visualization of DFC and GC organization in an MDA-MB-231 cell treated with SU056 at a concentration of 0.25 μM.

Video S5. Three-dimensional visualization of DFC and GC organization in an MDA-MB-231 cell treated with SU056 at a concentration of 2.0 μM.

Video S6. Three-dimensional visualization of DFC and GC organization in a control MCF-10A cell.

Video S7. Three-dimensional visualization of DFC and GC organization in an MCF-10A cell treated with 3% 1,6-hexanediol.

Video S8. Three-dimensional visualization of DFC and GC organization in an MCF-10A cell following washout of 1,6-hexanediol.

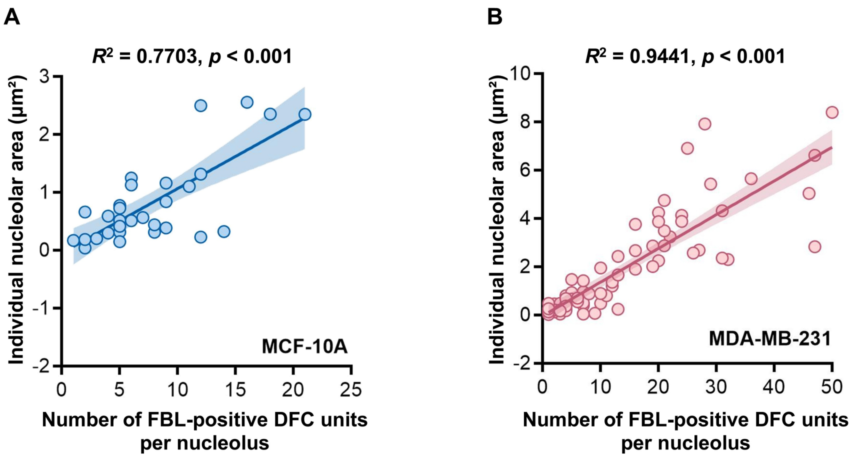


**Figure S1. Relationship between nucleolar area and the number of FBL-positive DFC units.**
(A–B) Scatter-density plots showing the relationship between the number of FBL-positive DFC units per nucleolus and individual nucleolar area in (A) MCF-10A and (B) MDA-MB-231 cells. Each point represents an individual nucleolus. Lines indicate linear regression fits. $R^2$ and $p$ values are shown for each analysis.
In both cell types, nucleolar area was positively associated with the number of FBL-positive DFC units.

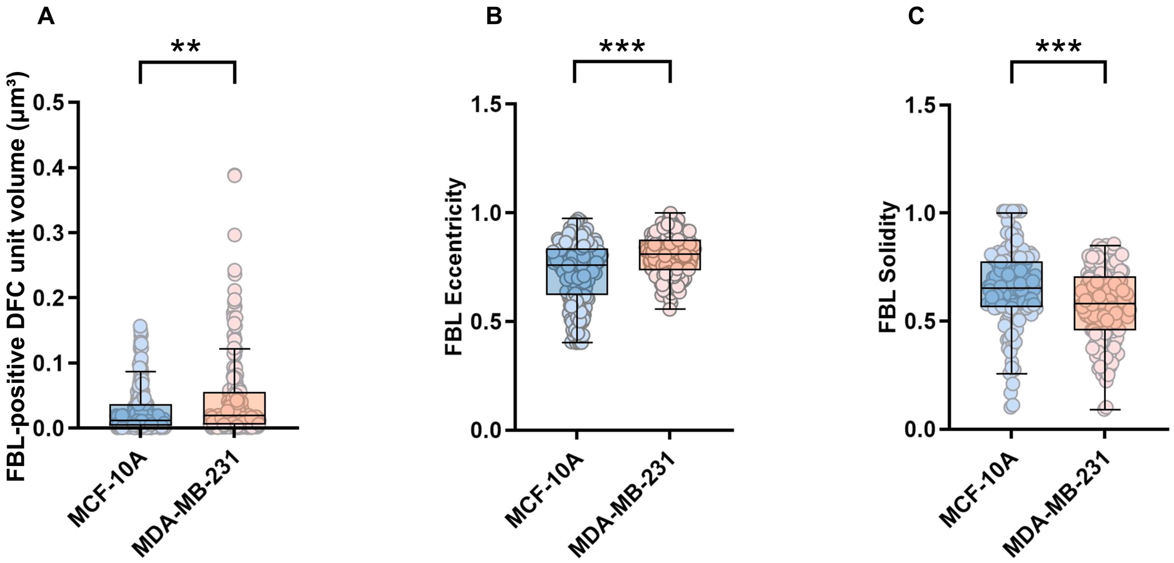


**Figure S2. Differences in the size and shape of individual FBL-positive DFC units in MCF-10A and MDA-MB-231 cells.**

(A) Estimated volume of individual FBL-positive DFC units, calculated from the projected area assuming spherical geometry. (B–C) Quantification of individual FBL-positive DFC unit shape, showing (B) eccentricity, with higher values indicating a more elongated shape, and (C) solidity, with lower values indicating greater boundary concavity.

A total of 770 FBL-positive DFC units from 100 nucleoli in 52 MCF-10A cells and 1,641 FBL-positive DFC units from 89 nucleoli in 54 MDA-MB-231 cells were analyzed across multiple experimental days. Data are presented as box-and-whisker plots with individual data points. Boxes indicate the median and interquartile range, and whiskers indicate the 1st–99th percentiles. Statistical significance was determined using Student's *t*-test. Cohen's d effect sizes were 0.55 for estimated volume, 0.60 for eccentricity, and 0.63 for solidity. ***p* < 0.01, ****p* < 0.001.

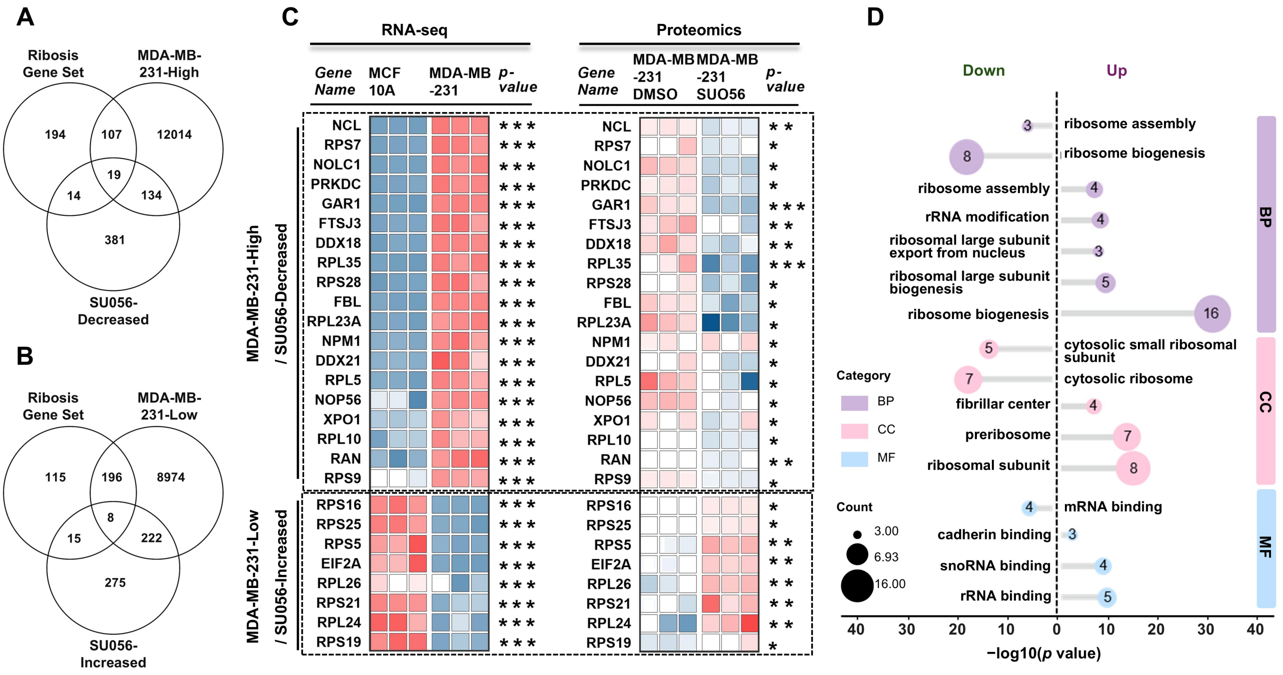


**Figure S3. Integrated transcriptomic and proteomic analysis identifies SU056-responsive ribosome-related factors in MDA-MB-231 cells.**

(A–B) Integrated identification of ribosome-related candidate genes showing opposing changes between MDA-MB-231 cells and SU056 treatment. (A) Intersection of the ribosome-related gene set, genes significantly upregulated in MDA-MB-231 cells relative to MCF-10A cells by RNA-seq, and proteins significantly decreased following SU056 treatment of MDA-MB-231 cells by mass spectrometry-based proteomics. 19 candidates were identified. (B) Intersection of the ribosome-related gene set, genes significantly downregulated in MDA-MB-231 cells relative to MCF-10A cells, and proteins significantly increased following SU056 treatment. 8 candidates were identified. (C) Heatmaps showing the relative expression profiles of candidates classified as Cancer-High/SU056-Decreased (top) and Cancer-Low/SU056-Increased (bottom). Cancer-High and Cancer-Low indicate genes with significantly higher or lower expression, respectively, in MDA-MB-231 cells relative to MCF-10A cells, whereas SU056-Decreased and SU056-Increased indicate proteins with significantly decreased or increased abundance, respectively, following SU056 treatment of MDA-MB-231 cells. Asterisks indicate statistical significance (*$p < 0.05$, **$p < 0.01$, ***$p < 0.001$). (D) Gene Ontology (GO) enrichment analysis of the identified candidates across Biological Process (BP), Cellular Component (CC), and Molecular Function (MF) categories. Circle size represents gene count, and the x-axis indicates −log10($p$ value).

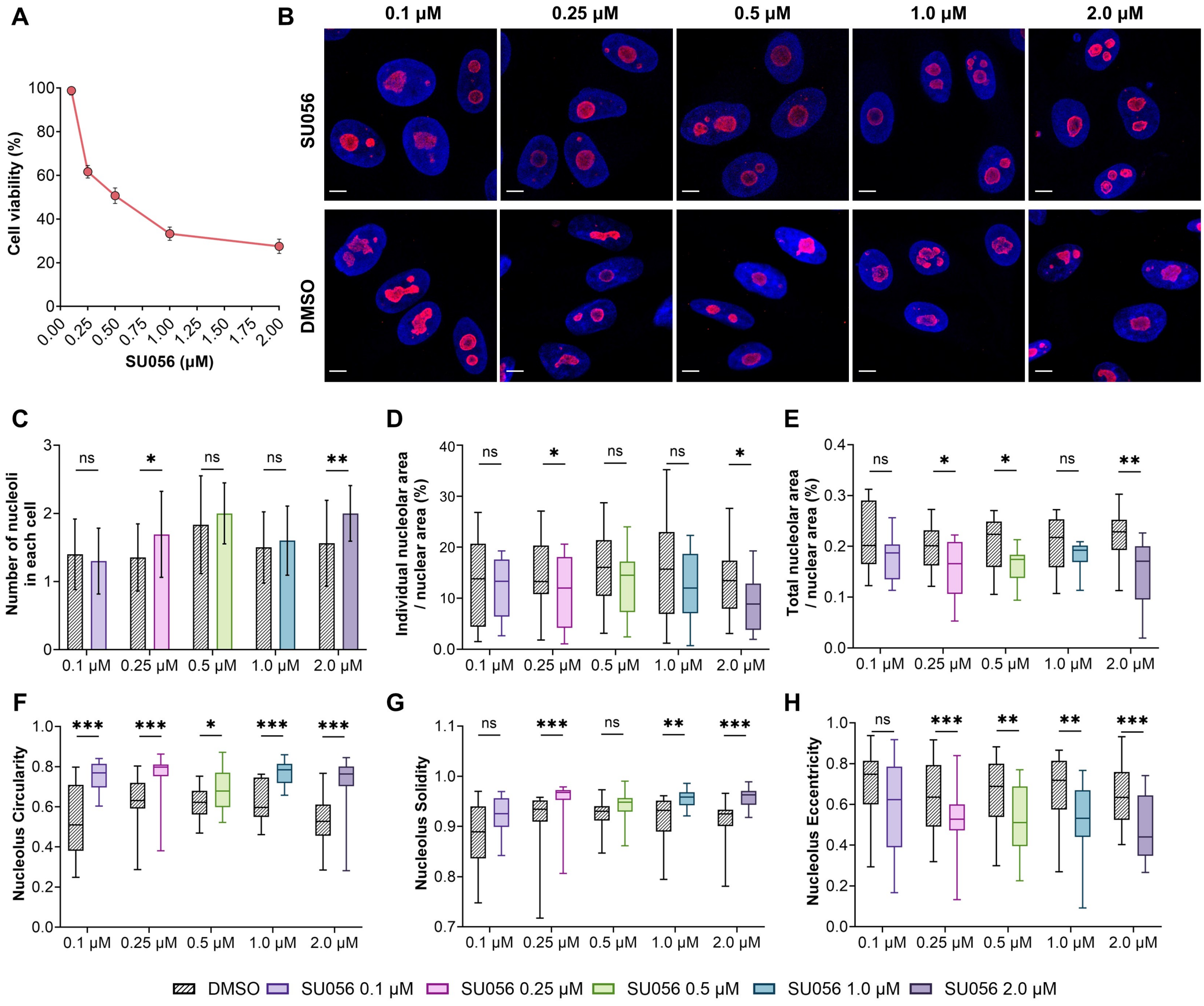


**Figure S4. Effects of increasing SU056 concentrations on cell viability and nucleolar morphology in MDA-MB-231 cells.**

(A) Cell viability of MDA-MB-231 cells treated with the indicated concentrations of SU056, expressed as a percentage of the control. (B) Representative fluorescence images showing nucleolar morphology in MDA-MB-231 cells treated with SU056 at concentrations of 0.1, 0.25, 0.5, 1.0, and 2.0 μM. NPM1 is shown in red and nuclei in blue. Scale bar = 5 μm. (C–H) Quantification of nucleolar morphological parameters across the SU056 concentration range: (C) number of nucleoli per cell, (D) individual nucleolar area relative to nuclear area, (E) total nucleolar area relative to nuclear area, (F) circularity, (G) solidity, and (H) eccentricity.

A total of 234, 136, 223, 147, 200, and 191 nucleoli from 137, 100, 141, 100, 111, and 101 cells were analyzed in the DMSO control, 0.1 μM SU056, 0.25 μM SU056, 0.5 μM SU056, 1.0 μM SU056, and 2.0 μM SU056 groups, respectively. Data are presented as mean ± SD in (A) and as box-and-whisker plots with individual data points in (C–H). Boxes indicate the median and interquartile range, and whiskers indicate the 1st–99th percentiles. Statistical comparisons were performed between SU056-treated and DMSO control groups. Statistical significance was determined using Student's *t*-test. *$p < 0.05$, **$p < 0.01$, ***$p < 0.001$; *ns*, not significant.

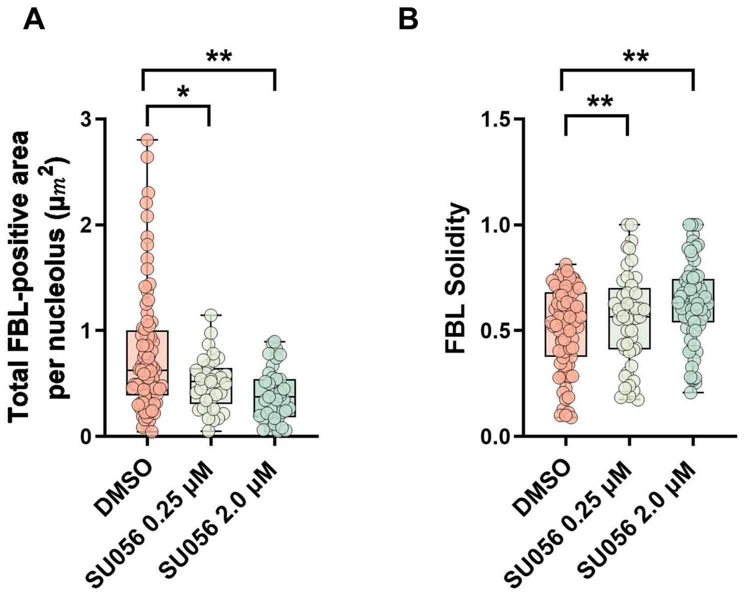


**Figure S5. Additional quantitative assessment of DFC organization following SU056 treatment in MDA-MB-231 cells.**

(A) Total FBL-positive area per nucleolus. (B) Solidity of individual FBL-positive DFC units. A total of 163, 322, and 370 nucleoli from 99, 167, and 114 cells and 2,028, 1,810, and 1,641 FBL-positive DFC units were analyzed in the DMSO group and in groups treated with SU056 at concentrations of 0.25 and 2.0 μM, respectively, across multiple experimental days. Data are presented as box-and-whisker plots overlaid with individual data points. Boxes indicate the median and interquartile range, and whiskers indicate the 1st–99th percentiles. Statistical significance was assessed using one-way ANOVA. *$p < 0.05$, **$p < 0.01$.

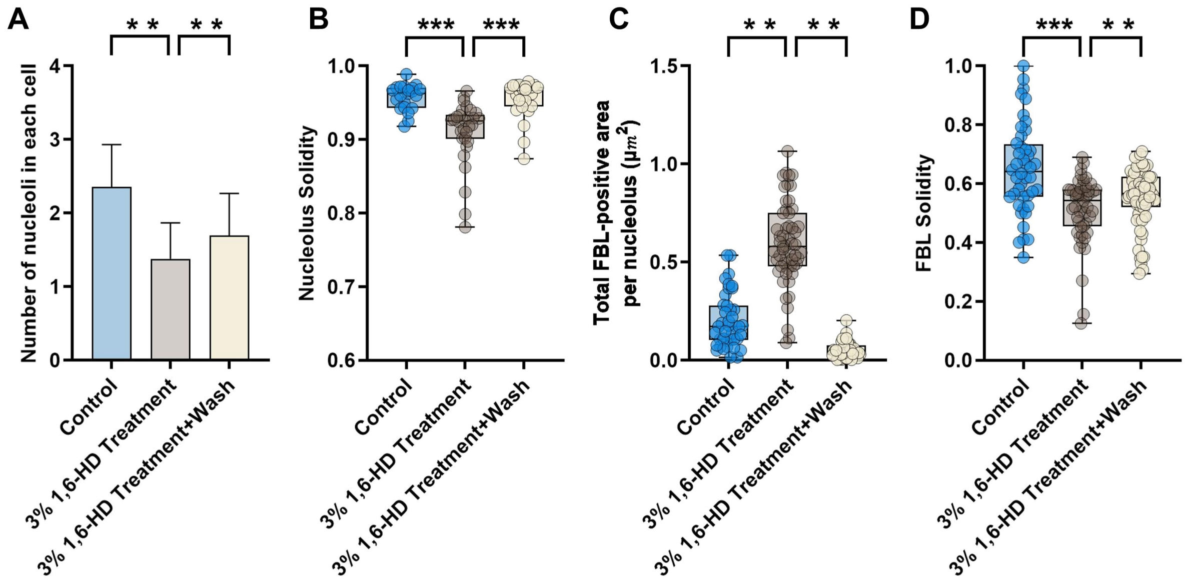


**Figure S6. Additional quantitative assessment of nucleolar morphology and DFC organization following 1,6-hexanediol treatment in MCF-10A cells.**

(A–B) Additional quantification of nucleolar morphology: (A) number of nucleoli per cell and (B) solidity. (C–D) Additional quantification of DFC organization: (C) total FBL-positive area per nucleolus and (D) solidity of individual FBL-positive DFC units.

A total of 54, 63, and 65 nucleoli from 27, 37, and 40 cells and 416, 945, and 1,170 FBL-positive DFC units were analyzed in the control, 3% 1,6-HD treatment, and washout groups, respectively, across multiple experimental days. Data are presented as box-and-whisker plots overlaid with individual data points. Boxes indicate the median and interquartile range, and whiskers indicate the 1st–99th percentiles. Statistical significance was assessed using one-way ANOVA. ***p* < 0.01, ****p* < 0.001.

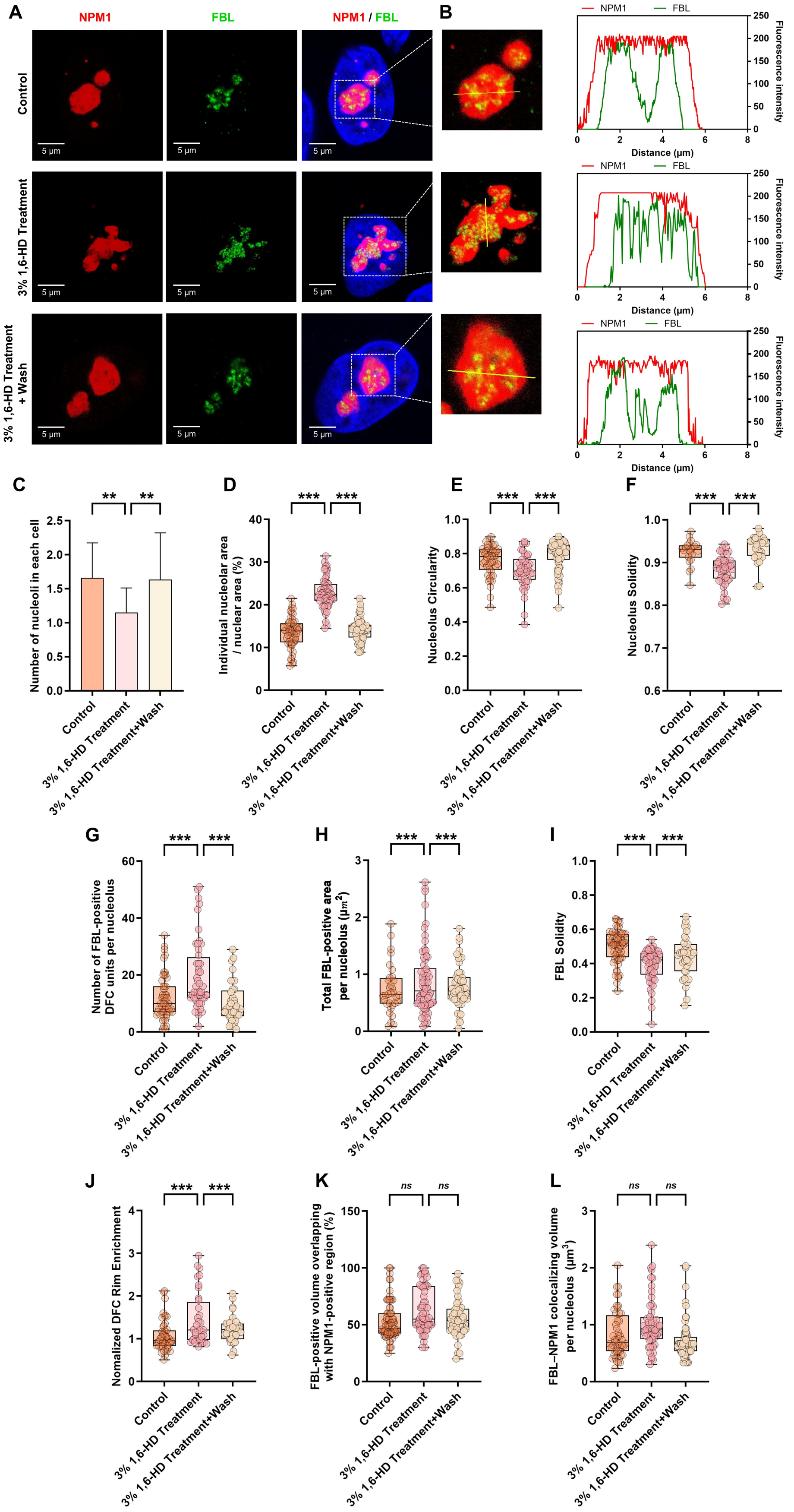

A
NPM1
FBL
NPM1 / FBL
B
Control
3% 1,6-HD Treatment
3% 1,6-HD Treatment + Wash
5 μm
NPM1
FBL
Distance (μm)
Fluorescence intensity
C
Number of nucleoli in each cell
D
Individual nucleolar area / nuclear area (%)
E
Nucleolus Circularity
F
Nucleolus Solidity
G
Number of FBL-positive DFC units per nucleolus
H
Total FBL-positive area per nucleolus (μm²)
I
FBL Solidity
J
Nomalized DFC Rim Enrichment
K
FBL-positive volume overlapping with NPM1-positive region (%)
L
FBL–NPM1 colocalizing volume per nucleolus (μm³)
Control
3% 1,6-HD Treatment
3% 1,6-HD Treatment+Wash
**
***
ns

**Figure S7. Effects of 1,6-hexanediol treatment on nucleolar morphology and DFC–GC organization in MDA-MB-231 cells.**

(A) Representative immunofluorescence images of NPM1 (red, GC) and FBL (green, DFC) in control MDA-MB-231 cells, MDA-MB-231 cells treated with 3% 1,6-hexanediol (1,6-HD), and cells following washout of 1,6-HD. Scale bar = 5 μm. (B) Representative line-profile analysis of NPM1 (red) and FBL (green) fluorescence intensities. Yellow lines in the magnified insets corresponding to the dashed boxes in (A) indicate the scan paths for the intensity profiles shown on the right. The x-axis represents distance along the scan line (μm), and the y-axis represents fluorescence intensity. (C–F) Quantification of nucleolar morphology: (C) number of nucleoli per cell, (D) individual nucleolar area relative to nuclear area, (E) circularity, and (F) solidity. (G–J) Quantification of DFC organization: (G) number of FBL-positive DFC units per nucleolus, (H) total FBL-positive area per nucleolus, (I) solidity of individual FBL-positive DFC units, and (J) normalized DFC rim enrichment. (K–L) Three-dimensional quantification of FBL–NPM1 spatial overlap: (K) percentage of FBL-positive volume overlapping with the NPM1-positive region per nucleolus and (L) total FBL–NPM1 colocalizing volume per nucleolus.

A total of 60, 67, and 70 nucleoli from 37, 42, and 43 cells and 1110, 1246, and 1330 FBL-positive DFC units were analyzed in the control, 3% 1,6-HD treatment, and washout groups, respectively, across multiple experimental days. Control cells received an equivalent volume of $H_2O$. Data are presented as box-and-whisker plots overlaid with individual data points. Boxes indicate the median and interquartile range, and whiskers indicate the 1st–99th percentiles. Statistical significance was assessed using one-way ANOVA. $^{**}p < 0.01$, $^{***}p < 0.001$; *ns*, not significant.

#### Video legends

**Video S1. Three-dimensional visualization of DFC and GC organization in an MCF-10A cell.**
3D reconstruction of a representative MCF-10A cell showing the spatial organization of FBL-positive DFC units and the NPM1-positive GC. The nucleus is shown in blue, NPM1 in red, and FBL in green. Regions of FBL–NPM1 overlap are shown in yellow.

**Video S2. Three-dimensional visualization of DFC and GC organization in an MDA-MB-231 cell.**
3D reconstruction of a representative MDA-MB-231 cell showing the spatial organization of FBL-positive DFC units and the NPM1-positive GC. The nucleus is shown in blue, NPM1 in red, and FBL in green. Regions of FBL–NPM1 overlap are shown in yellow.

**Video S3. Three-dimensional visualization of DFC and GC organization in a DMSO-treated MDA-MB-231 cell.**
3D reconstruction of a representative DMSO-treated MDA-MB-231 cell showing the spatial organization of FBL-positive DFC units and the NPM1-positive GC. The nucleus is shown in blue, NPM1 in red, and FBL in green. Regions of FBL–NPM1 overlap are shown in yellow.

**Video S4. Three-dimensional visualization of DFC and GC organization in an MDA-MB-231 cell treated with SU056 at a concentration of 0.25 µM.**
3D reconstruction of a representative MDA-MB-231 cell treated with SU056 at a concentration of 0.25 µM, showing the spatial organization of FBL-positive DFC units and the NPM1-positive GC. The nucleus is shown in blue, NPM1 in red, and FBL in green. Regions of FBL–NPM1 overlap are shown in yellow.

**Video S5. Three-dimensional visualization of DFC and GC organization in an MDA-MB-231 cell treated with SU056 at a concentration of 2.0 µM.**
3D reconstruction of a representative MDA-MB-231 cell treated with SU056 at a concentration of 2.0 µM, showing the spatial organization of FBL-positive DFC units and the NPM1-positive GC. The nucleus is shown in blue, NPM1 in red, and FBL in green. Regions of FBL–NPM1 overlap are shown in yellow.

**Video S6. Three-dimensional visualization of DFC and GC organization in a control MCF-10A cell.**
3D reconstruction of a representative control MCF-10A cell showing the spatial organization of FBL-positive DFC units and the NPM1-positive GC. The nucleus is shown in blue, NPM1 in red, and FBL in green. Regions of FBL–NPM1 overlap are shown in yellow.

**Video S7. Three-dimensional visualization of DFC and GC organization in an MCF-10A cell treated with 3% 1,6-hexanediol.**

3D reconstruction of a representative MCF-10A cell treated with 3% 1,6-hexanediol (1,6-HD), showing the spatial organization of FBL-positive DFC units and the NPM1-positive GC. The nucleus is shown in blue, NPM1 in red, and FBL in green. Regions of FBL–NPM1 overlap are shown in yellow.

**Video S8. Three-dimensional visualization of DFC and GC organization in an MCF-10A cell following washout of 1,6-hexanediol.**

3D reconstruction of a representative MCF-10A cell following washout of 3% 1,6-hexanediol (1,6-HD), showing the spatial organization of FBL-positive DFC units and the NPM1-positive GC. The nucleus is shown in blue, NPM1 in red, and FBL in green. Regions of FBL–NPM1 overlap are shown in yellow.
